# Discovery of late intermediates in methylenomycin biosynthesis active against drug-resistant Gram-positive bacterial pathogens

**DOI:** 10.1101/2025.05.11.653355

**Authors:** Christophe Corre, Gideon A. Idowu, Lijiang Song, Melanie E. Whitehead, Lona M. Alkhalaf, Gregory L. Challis

**Affiliations:** Department of Chemistry, University of Warwick, Coventry CV4 7AL, UK; School of Life Sciences, University of Warwick, Coventry CV4 7AL, UK; Department of Biochemistry and Molecular Biology, Biomedicine Discovery Institute, Monash University, Clayton, Victoria 3800, Australia; ARC Centre of Excellence for Innovations in Peptide and Protein Science, Monash University, Clayton Victoria 3800, Australia

## Abstract

The methylenomycins are highly functionalized cyclopentanone antibiotics produced by *Streptomyces coelicolor* A3(2). A biosynthetic pathway to the methylenomycins has been proposed based on sequence analysis of the proteins encoded by the methylenomycin biosynthetic gene cluster and incorporation of labelled precursors. However, the roles played by putative biosynthetic enzymes remain experimentally uninvestigated. Here, the biosynthetic functions of enzymes encoded by *mmyD, mmyO, mmyF* and *mmyE* were investigated by creating in-frame deletions in each gene and investigating the effect on methylenomycin production. No methylenomycin-related metabolites were produced by the *mmyD* mutant, consistent with the proposed role of MmyD in an early biosynthetic step. The production of methylenomycin A, but not methylenomycin C, was abolished in the *mmyF* and *mmyO* mutants, consistent with the corresponding enzymes catalyzing epoxidation of methylenomycin C, as previously proposed. Expression of *mmyF* and *mmyO* in a *S. coelicolor* M145 derivative engineered to express *mmr*, which confers methylenomycin resistance, enabled the resulting strain to convert methylenomycin C to methylenomycin A, confirming this hypothesis. A novel metabolite (pre-methylenomycin C), which readily cyclizes to form the corresponding butanolide (pre-methylenomycin C lactone), accumulated in the *mmyE* mutant, indicating the corresponding enzyme is involved in introducing the exomethylene group into methylenomycin C. Remarkably, both pre-methylenomycin C and its lactone precursor were one to two orders of magnitude more active against various Gram-positive bacteria, including antibiotic-resistant *Staphylococcus aureus* and *Enterococcus faecium* isolates, than methylenomycins A and C, providing a promising starting point for the development of novel antibiotics to combat antimicrobial resistance.

## INTRODUCTION

Methylenomycin A (**1**) is an unusual antibiotic produced by the model Actinomycete *Streptomyces coelicolor* A3(2) (Figure 1),^1-3^ with a wide spectrum of antibiotic activity, including against diverse Gram-positive bacteria and Gram-negative *Proteus* spp.^4^ The structurally related metabolites methylenomycin C (**2**), methylenomycin B (**3**) and xanthocidin (**4**) (Figure 1) have been isolated from several *Streptomyces* spp.^5-7^

**Figure 1.**
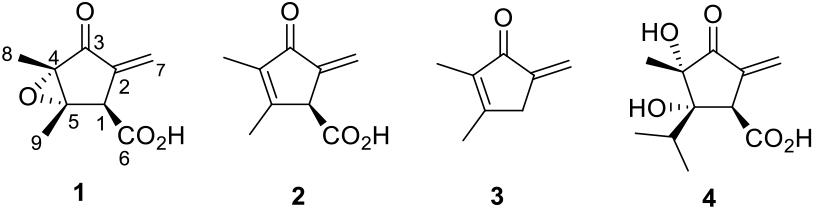
Structures of methylenomycin A (**1**), methylenomycin C (**2**), methylenomycin B (**3**), and xanthocidin (**4**).

Incorporation experiments with isotope-labelled precursors have identified the metabolic origin of the methylenomycins and shown that MmC (**2**) is a precursor of MmA (**1**).^8-10^ Furthermore, sequencing of the *S. coelicolor* A3(2) giant linear plasmid SCP1 identified the methylenomycin biosynthetic gene cluster.^11^ Based on sequence comparisons, thirteen proteins encoded by this gene cluster are proposed to play a role in methylenomycin biosynthesis (Figure 2 and Table S1). Three genes (*mmfL, mmfP* and *mmfH*) flanking the cluster of biosynthetic genes direct production of the methylenomycin furans (MMFs), a group of hormones that induce methylenomycin production.^10-14^ The nine carbon atoms of **1** and **2** derive from 2 molecules of acetate and a molecule of ribose,^8-10^ and MmyD shows 47% similarity to AvrD, which is proposed to catalyse the condensation of beta-ketoacyl thioesters with xyloluse in syringolide biosynthesis.^15^ Together, these observations led us to propose that MmyD catalyses the condensation of acetoacetyl-MmyA (assembled from acetyl- and malonyl-CoA by MmyC and a malonylltransferase borrowed from primary metabolism) with a pentulose, forming a butenolide intermediate that gets elaborated to **2**, which undergoes epoxidation catalysed by MmyF and MmyO to form **1** (Figure 2).^10^

**Figure 2.**
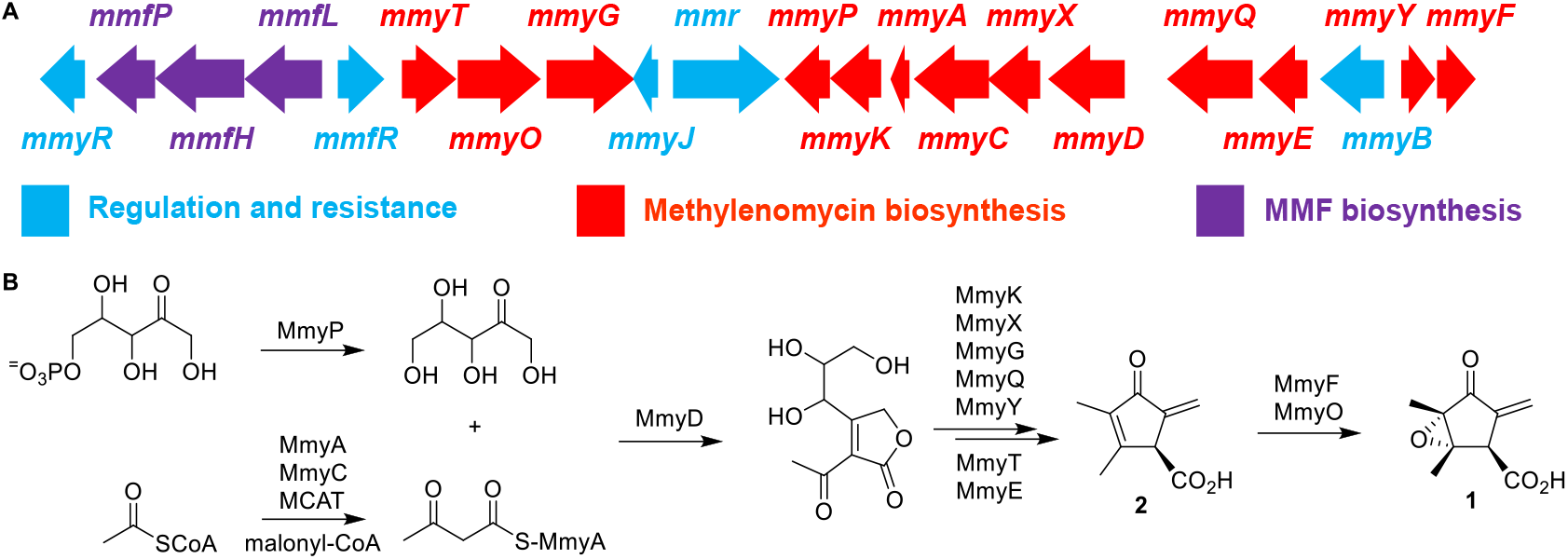
(**A**) Organization of the *S. coelicolor* methylenomycin biosynthetic gene cluster. The 13 genes implicated in methylenomycin biosynthesis are colored red. *mmyD, mmyE, mmyO*, and *mmyF* were deleted in this study; they encode enzymes with similarity to a putative butenolide synthase, flavin-dependent oxidoreductases, flavin-dependent monooxygenases, and a flavin reductase, respectively. (**B**) Proposed pathway for the biosynthesis of methylenomycins A (**1**) and C (**2**) in *S. coelicolor*. MCAT = malonyl-CoA acyl transferase from primary metabolism.

Although the pathway we propose for the biosynthesis of **1** is plausible, experimental evidence to support the proposed roles played by proteins encoded by the *mmy* gene cluster has hitherto been lacking. Here, we report in-frame deletion of four putative biosynthetic genes (*mmyD, mmyE, mmyO* and *mmyF*) in a cosmid containing the entire methylenomycin biosynthetic gene cluster, in parallel with deletion of *mmyR* to boost methylenomycin titres. The cosmids were integrated into the chromosome of *S. coelicolor* M145, which lacks SCP1, and the production of methylenomycin-related metabolites in each strain was analysed. The results of these experiments were consistent with proposed roles of MmyD, MmyF and MmyO in the methylenomycin biosynthesis, and further experiments confirmed that MmyF and MmyO catalyse the conversion of **2** to **1** using molecular oxygen. Two novel compounds, pre-methylenomycin C lactone **5** and pre-methylenomycin C **6**, accumulated in the *mmyE* mutant, suggest MmyE participates in the formation of the exomethylene group in **2**, via elimination of water from **6**. Surprisingly, **5** and **6** were significantly more active than **1** and **2** against various Gram-positive bacteria, including methicillin -resistant *Staphylcoccus aureus* (MRSA). The low MIC values for **5** against MRSA and *Enterococcus faecium* (1-2 µg/mL) suggest it may provide a promising starting point for the development of new antibiotics to tackle antimicrobial resistance.

## RESULTS AND DISCUSSION

### Construction and expression of mutated *mmy* gene clusters

Cosmid C73-787, containing the entire *mmy* gene cluster in addition to an integrative cassette that enables it to insert into the phage C31 *attB* site of *Streptomyces* chromosomes,^12,16^ was used to construct the desired mutants. *S. coelicolor* M145 containing this plasmid produces the methylenomycins.^12^ To investigate the biosynthetic roles of *mmyD, mmyE, mmyO* and *mmyF*, each of these genes was separately replaced in C73-787 by an apramycin resistance cassette using PCR-targeting (Figure S1 and Table S2).^17^ The resistance cassette was then excised leaving an 81 bp in-frame “scar” sequence between the start and stop codon of each gene.

MmyR represses expression of the *mmy* gene cluster.^13^ Consequently, deletion of *mmyR* boosts methylenomycin A titres in *S. coelicolor*.^13^ We therefore replaced *mmyR* with the apramycin resistance cassette in both the starting cosmid and each of the mutant cosmids (Figure S1). This also enabled selection for each construct upon transformation of *E. coli* ET12567/pUZ8002.

*S. coelicolor* M145 was transformed with each of the cosmids, and DNA integration was confirmed by PCR (Table S3 and Figures S2 and S3). The resultant strains were grown on supplemented agar minimal medium containing a higher phosphate concentration than typical *Streptomyces* fermentation media, such as R2YE. Increased phosphate has been shown to inhibit prodiginine and actinorhodin production in *S. coelicolor* without affecting the production of the methylenomycins.^2^ This increases precursor supply for the biosynthesis of the methylenomycins (and related metabolites), facilitating purification and spectroscopic analysis. The profile of metabolites produced by each strain was analysed by LC-MS.

### Deletion of *mmyD* abolishes methylenomycin production

As anticipated, *S. coelicolor* W89, containing the Δ*mmyR* derivative of C73-787, produced **1** and **2** in good titres (Figure 3). In contrast, the production of **1** and **2** was abolished in *S. coelicolor* W95, which contains the Δ*mmyD* / Δ*mmyR* derivative of C73-787, and no methylenomycin-related metabolites could be detected in this mutant (Figure 3). Transformation of W95 with an integrative plasmid containing *mmyD* under the control of the constitutive *ermE*\* promoter, resulting in *S. coelicolor* W118, restored methylenomycin production (Table S3 and Figure S4). This is consistent with our previous proposal that MmyD catalyses condensation of a pentulose with acetoacetyl-MmyA at an early stage in methylenomycin biosynthesis.^10^

**Figure 3.**
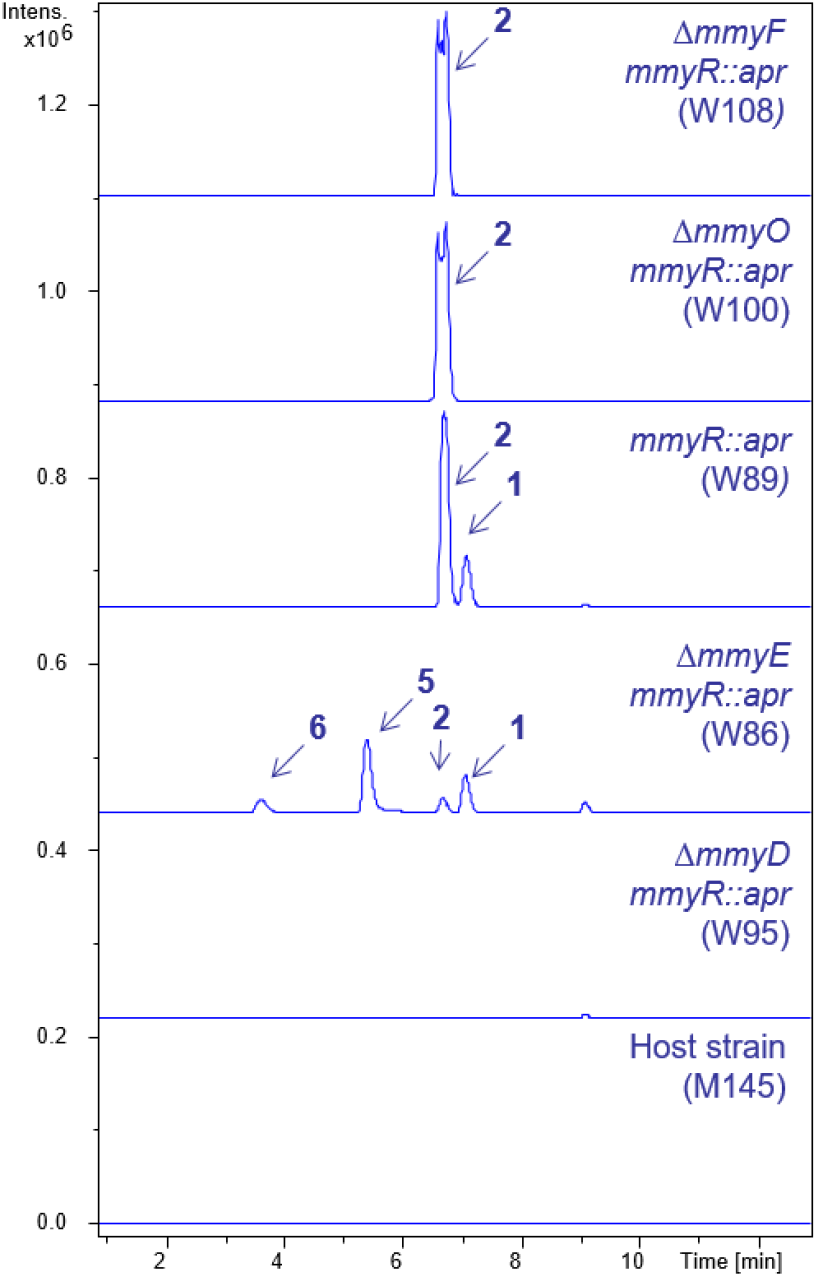
Extracted ion chromatograms at *m/z* = 139.0700 ± 0.005 (corresponding to the [M-CO_2_+H]^+^ ion of **1**) and *m/z* = 167.070 ± 0.005 (corresponding to the [M+H]^+^ ions of **2** and **5** and the [M-H_2_O+H]^+^ ion of **6**) from positive ion mode LC-MS analyses of neutral extracts (pH 5.5) of *S. coelicolor* M145 (bottom), W95 (second from bottom), W86 (third from bottom), W89 (third from top), W100 (second from top), and W108 (top).

### Novel metabolites accumulate in the *mmyE* mutant

The production of **1** and **2** was greatly reduced in *S. coelicolor* W86 (containing the Δ*mmyE* / Δ*mmyR* derivative of C73-787) and two new metabolites, absent from cultures of both *S. coelicolor* W89 and M145, were detected in neutral extracts (Figure 3). One exhibited a λ_max_ of 245 nm and had the molecular formula C_9_H_10_O_3_ (*m/z* = 167.0701 and 189.0518; calculated *m/z* for C_9_H_11_O_3_^+^ and C_9_H_10_O_3_Na^+^ = 167.0708 and 189.0521, respectively; Figure S5). The other had a λ_max_ of 236 nm and the molecular formula C_9_H_12_O_4_ (*m/z* = 167.0705, 185.0814, and 207.0632; calculated *m/z* for C_9_H_11_O_3_ ^+^, C_9_H_13_O _4_^+^ and C_9_H_12_O_4_Na^+^ = 167.0708, 185.0807, and 207.0629, respectively). In negative ion mode, the latter gave rise to an ion with *m/z* = 183.0650 corresponding to [M-H]^-^ for a species with the molecular formula C_9_H_12_O_4_.

In acidified (pH 3) extracts of *S. coelicolor* W86, the compound with molecular formula C_9_H_12_O_4_ could not be detected (Figure S6). The compound with molecular formula C_9_H_10_O_3_ was extracted from acidified large cultures using ethyl acetate and purified by preparative HPLC. Analysis of ^1^H, ^13^C, COSY, HSQC, HMBC, and ROESY NMR spectra (Figures S7-S12) showed this compound has structure **5** (Figure 4). The CD spectra of **1, 2** and **5** are very similar (Figure S13), leading us to conclude that all three compounds have identical absolute configurations.

**Figure 4.**
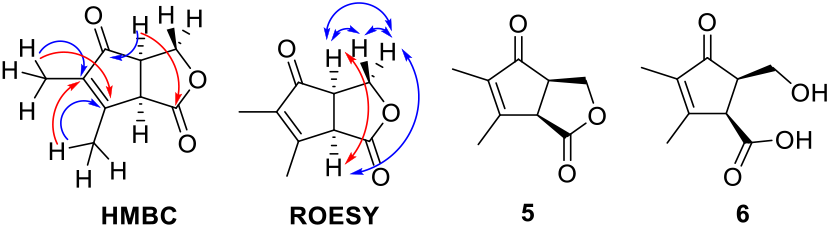
Summary of correlations observed (blue = strong, red = weak) in HMBC and ROESY spectra of pre-methylenomycin C lactone (**5**) and the structure of pre-methylenomycin C (**6**).

The compound with molecular formula C_9_H_12_O_4_ could not be isolated in sufficient quantity from neutral extracts of *S. coeliocolor* W86 to permit structure elucidation using NMR spectroscopy. However, **5** could be converted to this compound with NaOH in THF. ^1^H, ^13^C, COSY, HSQC, HMBC, and ROESY NMR spectroscopic analyses (Figures S14-S19) showed this compound has structure **6** (Figure 4).

Like other γ-hydroxy acids, accumulated **5** probably undergoes slow spontaneous lactonization to form **6**. As expected, this process is accelerated at lower pH. Dehydration of **6** would yield **2**. Thus, we propose the name pre-methylenomycin C for **6**. Stereoelectronic constraints prevent **5**, which we named pre-methylenomycin C lactone, from being converted directly to **2**. The protein encoded by *mmyT* shows sequence similarity to type II thioesterases and likely catalyses the conversion of **5** to **6**.

Our data indicate that MmyE catalyses the conversion of **6** to **2**. MmyE has 32% sequence identity to PlmM, a flavin-dependent enoylreductase from *Streptomyces sp*. HK803 involved in assembly of the cyclohexanecarboxyl-CoA starter unit for phoslactomycin biosynthesis.^18^ Interestingly, flavoenzymes usually catalyse redox reactions, but the conversion of **6** to **2** does not involve a net change in oxidation state. Other flavoenzymes have been reported to catalyse non-redox reactions.^19^ The bound flavin is proposed to play a purely structural role in these enzymes. The MmyE-catalysed conversion of **6** to **2** likely proceeds via an E1cB mechanism involving a basic residue in the active site that generates an enolate intermediate, which undergoes elimination of hydroxide promoted by an acidic active site residue (Scheme 1).

**Scheme 1.**
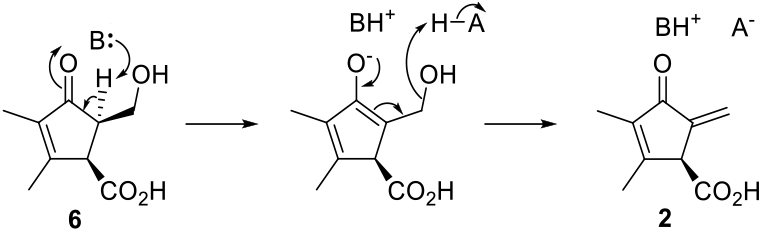
The MmyE-catalysed conversion of **6** to **2** likely proceeds via an E1cB mechanism involving a basic (B) and an acidic (A-H) active site residue.

### Methylenomycin A is not produced by the *mmyO* and *mmyF* mutants

The production of **1** but not **2** was abolished in *S. coelicolor* W100 and 108 (containing the Δ*mmyO* / Δ*mmyR* and Δ*mmyF* Δ*mmyR* derivatives of C73-787, respectively) (Figure 3). Two novel metabolites with the same molecular formula (C_9_H_12_O_3_; Figure S20) were detected in extracts of cultures grown for more than 3 days. Both compounds were purified from ethyl acetate extracts of acidified supernatants from 7-day cultures of *S. coelicolor* W108. Analysis of ^1^H, COSY, HSQC, and HMBC NMR spectra (Figures S21 – S28) showed the two compounds are diastereomers with structures **7** and **8**. These are likely shunt metabolites derived from unselective reduction of the exomethylene in **2** (Figure 5). They were named methylenomycins D1 (**7**) and D2 (**8**), respectively. Consistent with the shunt metabolite hypothesis, decreased levels of **2** were observed in cultures accumulating **7** and **8**, and in cultures grown for more than 72 hours **2** could no longer be detected. Comparison of the CD spectra for **7** and **8** with **6, 2**, and **1** suggested these metabolites all have the same absolute configuration at C2 (Figure S13).

**Figure 5.**
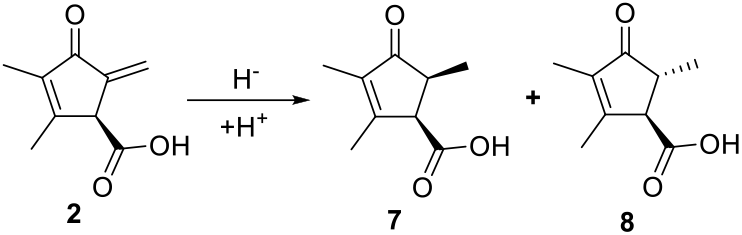
Structures of methylenomycins D1 (**7**) and D2 (**8**) isolated from *S. coelicolor* W108 proposed to derive from **2**.

### MmyO and MmyF convert methylenomycin C to A

MmyO has 43% sequence identity to LimB, a monooxygenase that catalyses FADH_2_–dependent epoxidation of the 1,2 double-bond of limonene in *Rhodococcus erythropolis*, using molecular oxygen as co-substrate.^20^ MmyF is show sequence similarity to NADPH-dependent flavin reductases. The inability of *S. coelicolor* W108 (containing *mmyO* but not *mmyF)* to produce **1** indicates there is a strict requirement for MmyF to supply FADH_2_ to MmyO (Figure 3). Attempts to overproduce MmyO and MmyF in *E. coli* were unsuccessful. Thus, we elected to co-express *mmyO* and *mmyF* in *S. coelicolor* M145. The two genes were amplified from C73_787 and cloned into pOSV556t under the control of the strong constitutive *ermE*\* promoter (Figure S29). The resulting vector was integrated into the chromosome of *S. coelicolor* M145 *via* intergeneric conjugation from *E. coli* ET12567 containing pUZ8002, resulting in *S. coelicolor* W110.

Purified **2** was fed to cultures of *S. coelicolor* W110 and M145. However, both died, presumably because these strains are sensitive to **2**. To circumvent this problem, *mmr*, which has been shown to confer methylenomycin resistance, was amplified from C73_787 (Table S4) and cloned into the multi-copy plasmid pIJ86 under the control of the *ermE*\* promoter (Figure S30). The resulting construct was introduced into *S. coelicolor* M145 and W110 via intergenic conjugation from *E. coli*, resulting in *S. coelicolor* W301 and W302, respectively. To examine methylenomycin resistance, filter paper discs saturated with **1** were placed on plates inoculated separately with *S. coelicolor* M145, W110, W301, and W302. After 5 days of incubation, a sizeable zone of growth inhibition was observed for *S. coelicolor* M145 and W110, whereas the growth of *S. coelicolor* W301 and W302 was unaffected (Figure S30). This confirmed that *S. coelicolor* W301 and W302 are resistant to **1** the putative product of MmyO / MmyF catalysis.

*S. coelicolor* W301 and W302 were grown on AlaMM (pH 5.0) agar medium for 2 days and **2** was added. After 3 days further incubation, the agar was extracted with methanol and the concentrated extracts were analysed using LC-ESI-Q-ToF-MS. This showed that **1** was produced by *S. coelicolor* W301 (containing *mmyO, mmyF* and *mmr*) but not W302 (containing just *mmr*), confirming MmyO and MmyF togther catalyse the conversion of **2** to **1** (Figure 6).

**Figure 6.**
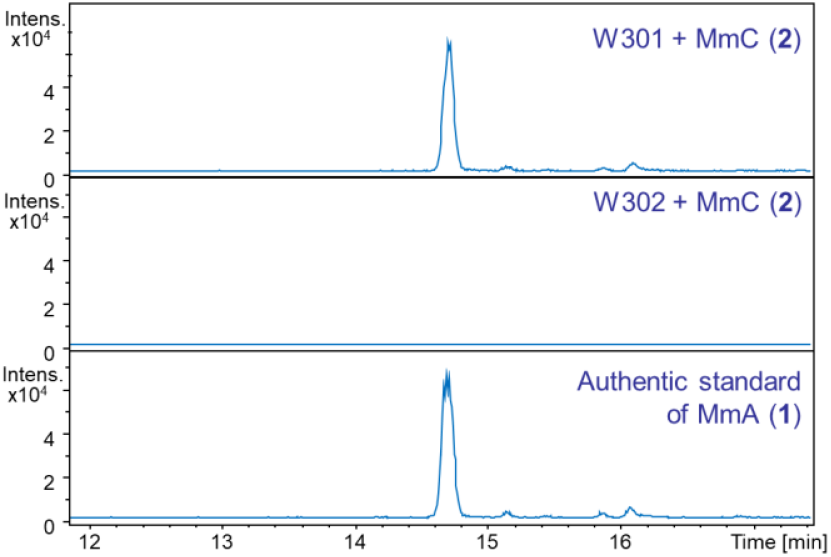
Extracted ion chromatograms (EICs) at *m/z* = 183.065 ±0.005 corresponding to [M+H]^+^ for **1** from UHPLC-ESI-Q-ToF-MS analyses of methanol extracts from *S. coelicolor* W301 (top panel) and W302 (middle panel) fed with **2**. The EIC at *m/z* = 183.065 ±0.005 for an authentic standard of **1** purified from *S. coelicolor* W89 is shown in the bottom panel.

To further characterise the reaction catalysed by MmyO / MmyF, we investigated the origin of the oxygen atom in the epoxide group of **1**, which we hypothesized derives from molecular oxygen. *S. coelicolor* W89 was grown under an ^18^O_2_ atmosphere and UHPLC-ESI-Q-ToF-MS analysis of culture extracts revealed just over 50% incorporation of a single ^18^O atom into **1**, whereas no ^18^O incorporation into **2** was observed (Figure S31). The lower than 100% incorporation of ^18^O into **1** is likely due to incomplete exclusion of air from the agar cultures. Because **1** is derived from **2** by converting a C=C into an epoxide and no ^18^O_2_ is incorporated into **2**, the epoxide oxygen of **1** must derive from molecular oxygen.

The conversion of **2** to **1** likely proceeds via the reaction of MmyO-bound FADH_2_, generated by MmyF-mediated reduction of FAD, with O_2_ (Scheme 2). The resulting C4a-peroxyflavin can add to C5 of **2**, creating an enolate intermediate that collapses via nucleophilic attack of C4 on the peroxide. An analogous mechanism has been proposed for the epoxidation of electron deficient double bonds by other flavoenzymes.^21^

### Substrate tolerance of MmyO

To investigate whether MmyO has a broad substrate scope, we fed **5, 6, 7**, and **8**, all of which have the same cyclopentenone core as **2**, to *S. coelicolor* W301 and W302 (as a negative control). LC-MS analysis of culture extracts (Figures S32 and S33) showed that **6** was the only compound turned over by *S. coelicolor* W301 to a monooxygenated product. Re-examination of

*S. coelicolor* W89 culture extracts showed that this strain produces the same monooxygenated derivative of **6**. Thus, this metabolite was purified by HPLC from large scale cultures of W89. HRMS showed this compound has the molecular formula C_9_H_12_O_5_ (*m/z* calculated for [C_9_H_12_O_5_Na]^+^ = 223.0576, measured *m/z* = 223.0577; Figure S34) consistent with structure **9**, the anticipated product of the MmyO-catalysed epoxidation of **6**. Surprisingly, ^1^H, COSY, HSQC and HMBC NMR spectroscopic analyses (Figures S35 – S38) led us to conclude that this compound has structure **10**, which can be formed from **9** via an intramolecular rearrangement (Scheme 3). Simple commercially available analogues of **2**, such as cylopentenone and cyclohexanone, were also fed to *S. coelicolor* W301, but no monooxygenated derivatives could be detected.

**Scheme 2.**
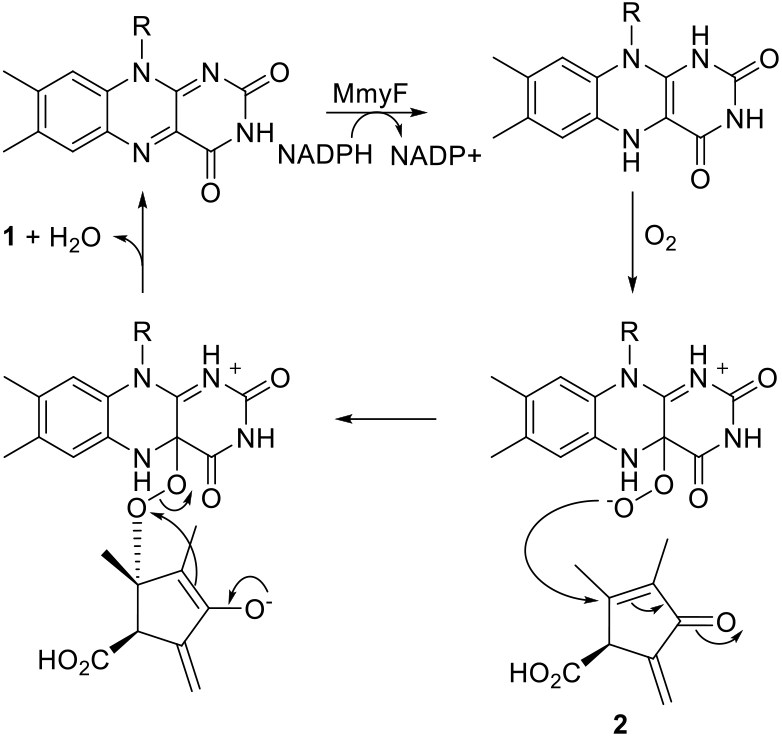
Proposed mechanism for the conversion of **2** to **1**.

**Scheme 3.**
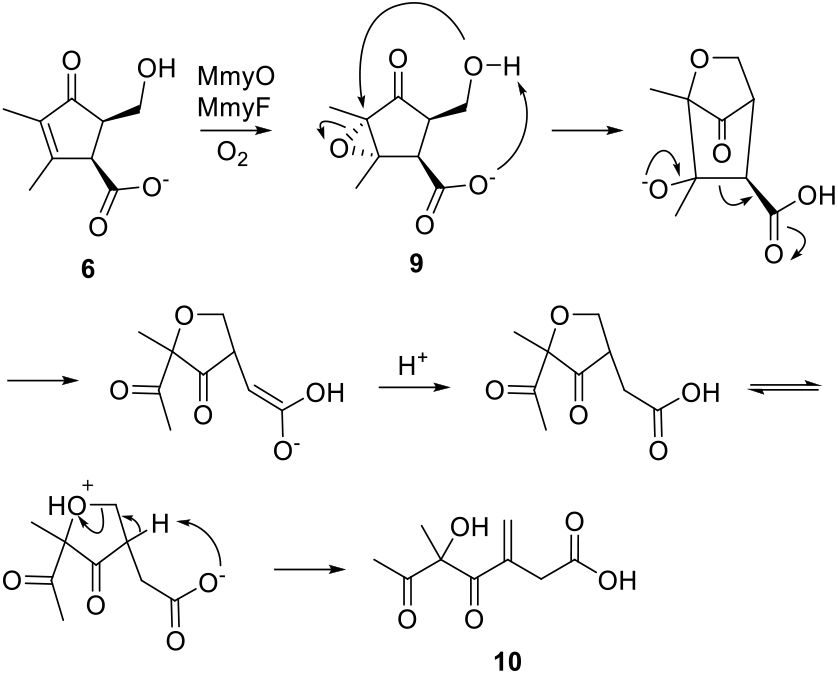
Proposed mechanism for formation of **10** from **9**, the presumed product of MmyO-catalysed epoxidation of **6**.

### Antimicrobial activity of intermediates, products, and shunt metabolites

The antimicrobial activity of **5, 6, 8**, and **9** against diverse Gram-positive bacteria, Gram-negative bacteria, and *Candida albicans* was compared to that of **1** and **2**. Minimum inhibitory concentrations (MICs) and minimum bactericidal concentrations (MBCs) for active compounds were determined using standard procedures.^21-23^

None of the compounds were active against Gram-negative bacteria (Table 1), presumably because they are unable to penetrate the outer membrane. **8** and **9** had no detectable activity against Gram-positive bacteria or *C. albicans* up to a maximum concentration of 512 μg/ml (Table 1). This is consistent with a previous report that reduction of the exomethylene group in methylenomycin A results in no biological activity at concentrations up to 400 µg/ml.^4c^ The exomethylene group in **1** and **2** thus appears to be a key pharmacophore.

**Table 1.**
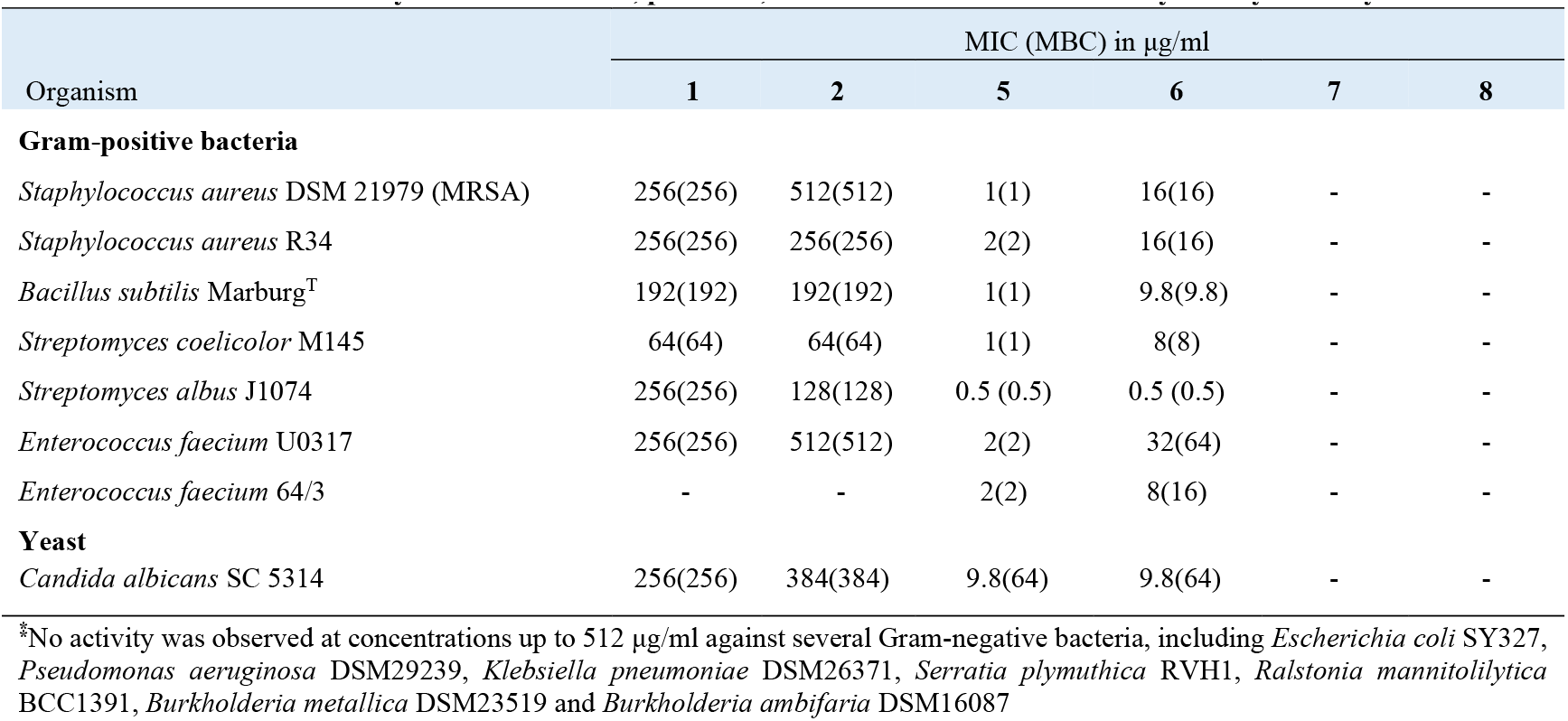
Antimicrobial activity^⁑^ of intermediates, products, and shunt metabolites in methylenomycin biosynthesis.

| Organism | MIC (MBC) in µg/ml |  |  |  |  |  |
| --- | --- | --- | --- | --- | --- | --- |
|  | 1 | 2 | 5 | 6 | 7 | 8 |
| **Gram-positive bacteria** |  |  |  |  |  |  |
| *Staphylococcus aureus* DSM 21979 (MRSA) | 256(256) | 512(512) | 1(1) | 16(16) | - | - |
| *Staphylococcus aureus* R34 | 256(256) | 256(256) | 2(2) | 16(16) | - | - |
| *Bacillus subtilis* Marburg<sup>T</sup> | 192(192) | 192(192) | 1(1) | 9.8(9.8) | - | - |
| *Streptomyces coelicolor* M145 | 64(64) | 64(64) | 1(1) | 8(8) | - | - |
| *Streptomyces albus* J1074 | 256(256) | 128(128) | 0.5 (0.5) | 0.5 (0.5) | - | - |
| *Enterococcus faecium* U0317 | 256(256) | 512(512) | 2(2) | 32(64) | - | - |
| *Enterococcus faecium* 64/3 | - | - | 2(2) | 8(16) | - | - |
| **Yeast** |  |  |  |  |  |  |
| *Candida albicans* SC 5314 | 256(256) | 384(384) | 9.8(64) | 9.8(64) | - | - |

\*No activity was observed at concentrations up to 512 µg/ml against several Gram-negative bacteria, including *Escherichia coli* SY327, *Pseudomonas aeruginosa* DSM29239, *Klebsiella pneumoniae* DSM26371, *Serratia plymuthica* RVH1, *Ralstonia mannitolilytica* BCC1391, *Burkholderia metallica* DSM23519 and *Burkholderia ambifaria* DSM16087

In our hands, **2**, which was previously reported to be inactive against *B. subtilis* subtilis,^5^ had similarly modest levels of activity as **1** against Gram-positive bacteria and *C. albicans* (Table 1). Surprisingly, **6** was an order of magnitude more active across the board than **1** and **2** (Table 1). Even more surprisingly, **5** was two orders of magnitude more active than **1** and **2** against all Gram-positive bacteria tested and had a similar level of activity to **6** against *C. albicans* (Table 1).

The MIC values for **5** of 1 and 2 µg/ml against the clinical isolates *Staphylococcus aureus* DSM 21979 and *Enterococcus faecium* U0317, respectively, are particularly noteworthy. *S. aureus* DSM 21979 is resistant to methicillin and aminoglycosides, whereas *E. faecium* U0317 is resistant to multiple classes on antibiotic, including chloramphenicol, macrolides, aminoglycosides, β-lactams, and tetracyclines. *E. faecium* U0317 and 64/3 are both susceptible to vancomycin, an antibiotic widely used for the treatment of enterococcal infections, with MBCs of 64 and 128 µg/ml, respectively.^24-26^ Strikingly, **5** has an MBC of 2 µg/ml against both these strains.

The acquisition of vancomycin resistance is a significant problem for the treatment of *E. faecium* infections.^24-26^ To investigate whether *E. faecium* 64/3 can evolve resistance to **5**, using vancomycin as a control, it was subjected to sequential passage through increasing concentrations of the antibiotics over a period of 28 consecutive days. This resulted in mutants with an 8-fold increase in MIC for vancomycin (from 4 µg/ml to 32 µg/ml), whereas the MIC for **5** remained unchanged (2 µg/ml). Thus, *E. faecium* appears unable to easily develop resistance to **5**.

The fact that **5** displays excellent antimicrobial activity, despite lacking the exomethylene group responsible for the weaker activity of **1** and **2**, demonstrates it employs a different pharmacophore. γ-Hydroxy acids are known to spontaneously lactonize, although this is often slow under neutral conditions. Partial lactonization of **5** to form **6** during antimicrobial activity assays could explain the observation of lower but still significant activity for the latter. These considerations suggest the γ-butyrolactone may be the pharmacophore in **6**.

## Conclusions

In this study, we employed a genetic approach to investigate the biosynthetic role of putative enzymes encoded by four genes in the *S. coelicolor* methylenomycin BGC. Abolition of the production of all methylenomycin-related metabolites in the *mmyD* mutant is consistent with our previous proposal that this gene encodes an AvrD-like enzyme responsible for condensing **-** no activity observed at concentrations up to 512 μg/ml A a β-keto-ACP thioester with a pentulose at an early stage in methylenomycin biosynthesis (Scheme 4). The structures of two novel methylenomycin-related metabolites, pre-methylenomycin C lactone (**5**) and pre-methylenomycin C (**6**), accumalated in the *mmyE* mutant suggest they are biosynthetic intermediates. We propose that the lactone in **5**, which is formed by the MmyD-catalysed reaction, is carried through subsequent steps catalysed by MmyG, MmyK, MmyQ, MmyY, and MmyX, resulting in assembly of the cyclopentanone (Scheme 4). Hydrolysis of the lactone in **5** by MmyT, which shows sequence similarity to type II thioesterases, would yield **6** (Scheme 4). Conversion of **6** to methylenomyicn C **2** is proposed to be catalysed by MmyE, which appears to be a redox-inactive flavoenzyme (Scheme 4). The observation that small amounts of **1** and **2** are still produced by the *mmyE* mutant suggests that another enzyme encoded by a gene outside the methylenomycin BGC can also catalyse the conversion of **6** to **2**, albeit inefficiently. A series of experiments demonstrate that MmyO and MmyF together catalyse the finals step methylenomycin A (**1**) biosynthesis – epoxidation of the tetrasubstituted double bond in **2**. MmyO, which we propose is a flavin-dependent monooxygenase supplied with reduced flavin by the reductase MmyF, appears to have a narrow substrate tolerance, although it can epoxidize **6** to make an unstable product **9** that undergoes a spectacular series of rearrangements to form **10**. Finally, two novel methylenomycin-related metabolites **7** and **8** were observed to accumulate in *mmyF* and *mmyO* mutants, in addition to the parent strain when grown for extended periods. These appear to result from non-specific reduction of the exomethylene group in **2** (Scheme 4). Overall, these studies afford considerable additional insight into methylenomycin biosynthesis, providing several testable new hypotheses, and indicating future efforts should focus on the mid-pathway roles play by MmyG, MmyK, MmyQ, MmyX, and MmyY.

**Scheme 4.**
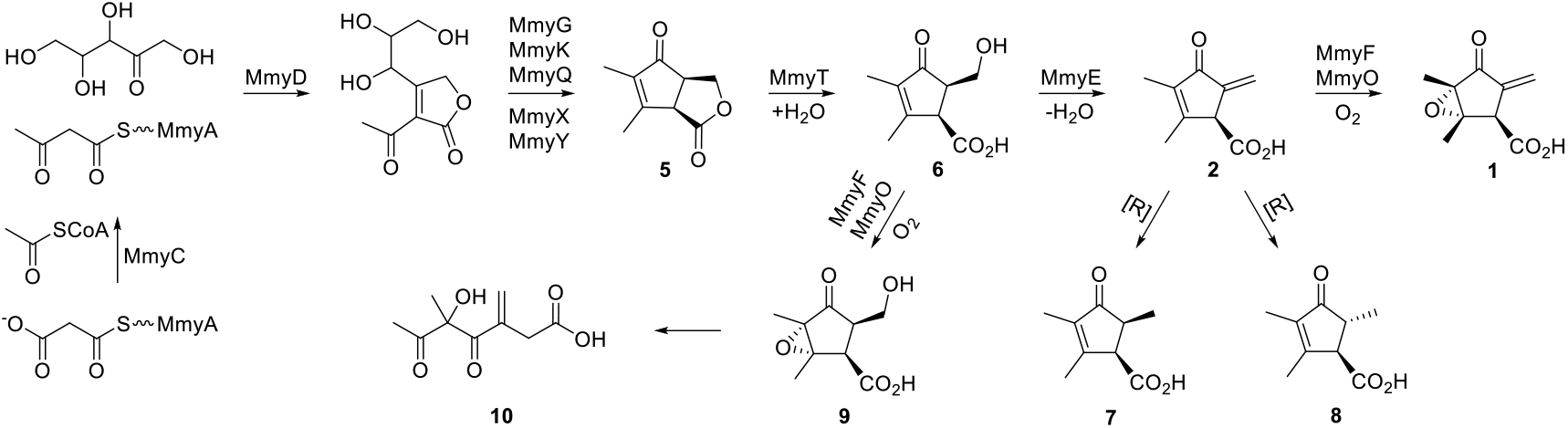
Revised pathway for methylenomycin biosynthesis in *S. coelicolor* A 3(2), based on the evidenced roles of MmyD, MmyE, MmyF and MmyO and on the isolation and structural characterization of metabolites 5, 6, 7 and 8.

Antimicrobial activity assays of the novel methylenomycin-related metabolites discovered in this work showed that the key biosynthetic intermediate pre-methylenomycin C lactone (**5**) is two orders of magnitude more active against diverse Gram-positive bacteria than methylenomycin A (**1**), the ultimate metabolic product. This suggests the methylenomycin BGC may initially have evolved to make the potent antibiotic **5**, with subsequent acquisition of the *mmyT, mmyE, mmyO*, and *mmyF* genes diverting the pathway first to **2** and then **1**, which have an alternative biological function. Identification and testing of intermediates in the biosynthesis of other metabolites with weak or no antimicrobial activity may therefore provide a fruitful new approach to antibiotic discovery. The activity of **5** against drug-resistant clinical isolates of *S. aureus* and *E. faecium*, coupled with its relatively simple structure and the apparent difficulty of evolving resistance to this compound in the latter, are all notable. It suggests **5** may provide a useful starting point for the development of novel antibiotics to tackle infections caused by multidrug-resistant Gram-positive bacteria. To this end, an expedient and versatile synthesis of **5** has been developed in collaboration with the Lupton group.^27^ This should enable the creation of diverse analogues that can be used to probe the structure-activity relationship and mechanism of action.

## ASSOCIATED CONTENT

### Supporting Information

Description of methods used, and supplementary figures and tables. This material is available free of charge via the Internet at http://pubs.acs.org..

## AUTHOR INFORMATION

### Author Contributions

CC and GAI contributed equally.

### Notes

The authors declare the following competing financial interests: G.L.C. is a Non-Executive Director, shareholder, and paid consultant of Erebagen Ltd.

## ACKNOWLEDGMENTS

We thank Dr Nikola Chmel for assistance with measuring CD spectra. This work was supported by a grant from the U.K. BBSRC (grant ref. BB/E008003/1) and the EU integrated project Actinogen (Contract No. 005224). GAI was the recipient of a Chancellor’s International Scholarship from the University of Warwick. GLC and CC were the recipients of a Wolfson Research Merit Award and a University Research Fellowship from the Royal Society (grant nos. WM130033 and UF090255).

