## Supporting Information for "Discovery of late intermediates in methylenomycin biosynthesis active against drug-resistant Gram-positive bacterial pathogens"

**Experimental procedures**

**Culture conditions for production of methylenomycins.** The *Streptomyces* strains were grown on a modified SMMS agar medium,[1] containing 5mM each of $NaH_2PO_4$ and $K_2HPO_4$. After incubation at 30 °C for 72 h, the plates were frozen overnight. After defrosting, the agar and mycelia were filtered with cotton wool by centrifugation for 5 mins at 2000 rpm. The resulting aqueous sample was analysed by LC-MS or used to purify methylenomycin related metabolites.

**Analytical methods.** LC-MS analyses were carried out using a reverse phase column (Agilent C18, 150 x 4.6 mm, 5 µm) connected to an Agilent 1100 HPLC instrument. The outflow was routed to a Bruker (MaXis$^{TM}$ Impact) High Resolution-Mass Spectrometer (HR-MS) fitted with an electrospray ionisation (ESI) source operating in positive or negative mode.

Methylenomycin related metabolites were separated using the following HPLC procedure from 20 uL of crude extract. Solvent A = $H_2O$ (0.1 % formic acid) Solvent B = MeOH (0.1 % formic acid). A:B 5:25 (0-5 min), 5:25 to 63:37 (5-21 min), 63:37 -45:55 (21-30 min), 45:55 -0:100 (30-35 min), 0:100 (35-40 min), 0:100 -75:25 (40-45 min), 75:25 (45-55 min).

**Purification and characterisation of pre-methylenomycin C lactone (5)**

SMMS agar (16 x 50 mL), was inoculated with spores of *S. coelicolor* W86. After incubation for 5 days at 30 ºC, cultures were combined, extracted with ethyl acetate (800 mL) and dried over $MgSO_4$. The solvent was removed *in vacuo* and the residue redissolved in water/methanol (1:1, 2 mL). Pre-MmCl was purified by preparative HPLC. Solvent A = $H_2O$ (0.1 % formic acid) Solvent B = MeOH (0.1 % formic acid). A:B 95:5 (0-5 min), 95:5 to 0:100 (5-25 min). Flow rate 20 mLmin$^{-1}$

The collected fractions were combined, $CH_3OH$ was removed *in vacuo* and the remaining aqueous fraction was extracted with ethyl acetate (2 x 50 mL). The ethyl acetate was dried over $MgSO_4$, filtered and removed *in vacuo*. NMR spectroscopy was conducted on a Bruker AV700 spectrometer equipped with a TCI cryoprobe.

**$\delta_H$** (700 MHz, $CDCl_3$), 1.73 (3H, s, C8), 2.25 (3H, s, C9), 3.29 (1H, dt, J 7.30, 3.30, H2), 3.63 (1H, d, J 7.30, H1), 4.40 (1H, dd, J 3.30, 9.70, H7b), 4.52 (1H, t, J 9.70, H7a) **$\delta_C$** (175 MHz, $CDCl_3$), 8.3 (C8),

15.5 (C9), 45.1 (C2), 48.8 (C1), 68.3 (C7), 137.3 (C5), 165.6 (C4), 173.3 (C6), 206.8 (C3), HR-ESI-MS *m/z* 167.0701 $[M + H]^+$ (Calculated for $[C_9H_{11}O_3]^+$: 167.0708).

**Synthesis and characterisation of pre-methylenomycin C (6)**

**5** (1 mg) was resuspended in THF (250 µL) and sodium hydroxide (10 mM, 250 µL) was added. The mixture was stirred at room temperature for 6 hours. The product was dried *in vacuo*

$\delta_H$ (700 MHz, $D_2O$), 1.71 (3H, s, H8), 2.09 (3H, s, H9), 2.88 (1H, q, J 6.70, 5.90, H2), 3.76 (1H, d, J 6.70, H1), 3.79 (1H, dd, J 5.90, 12.70, H7b), 3.89 (1H, dd, J 12.70, H7a), $\delta_C$ (176 MHz, $D_2O$), 7.2 (C8), 15.3 (C9), 50.3 (C2), 56.5 (C1), 59.9 (C7), 136.8 (C5), 173.2 (C4), 213.0 (C3), signal for C6 was not observed due to low intensity; HRMS Calculated for $[C_9H_{13}O_4]^+$: 185.0814, observed: 185.0807.

**Purification and characterisation of methylenomycin D1 (7) and methylenomycin D2 (8)**

SMMS agar (12 x 50 mL) was inoculated with spores of *S. coelicolor* W108 and incubated for 7 days at 30 ºC. The plates were combined, acidified to pH 3 and extracted with 700ml of ethyl acetate. The ethyl acetate was dried over magnesium sulphate and removed *in vacuo*. MmD1 (**7**) and MmD2 (**8**) were first copurified by silica chromatography (toluene-acetic acid 9:1). The mixture of diastereoisomers was then separated by preparative HPLC using the elution conditions described for **5**, except $CH_3CN$ was used in place of $CH_3OH$ to give **9** (2 mg) and **10** (4mg).

Methylenomycin D1 (**7**): $^1$H NMR (700 MHz, $CDCl_3$): $\delta$ 1.20 (3H, d, 3H's on C7, $J$ = 7.5 Hz), 1.76 (3H, s, 3H's on C8), 2.07 (3H, s, 3H's on C9), 2.72 (1H, m, H2), 3.77 (1H, d, H1, $J$ = 7.3 Hz) ppm. $^{13}$C NMR (175 MHz, $CDCl_3$): $\delta$ 8.3 (C8), 11.9 (C7), 15.8 (C9), 41.9 (C2), 53.2 (C1), 137.8 (C5), 163.3 (C4), 208.5 (C3) ppm. HR-ESI-MS *m/z* = 169.0860 $[M + H]^+$ (Calculated $[C_9H_{13}O_3]^+$: 169.0860).

Methylenomycin D2 (**8**): $^1$H NMR (700 MHz, $CDCl_3$): $\delta$ 1.27 (3H, d, 3H's on C7, $J$ = 7.4 Hz), 1.75 (3H, s, 3H's on C8), 2.09 (3H, s, 3H's on C9), 2.65 (1H, m, H2), 3.21 (1H, s, H1) ppm. $^{13}$C NMR (175 MHz, $CDCl_3$): $\delta$ 8.4 (C8), 15.5 (C9), 15.6 (C7), 44.3 (C2), 56.2 (C1), 137.3 (C5), 162.6 (C4), 208.5 (C3) ppm. HR-ESI-MS *m/z* = 169.0861 $[M + H]^+$ (Calculated $[C_9H_{13}O_3]^+$: 169.0860).

**Purification of methylenomycin C (2)**

Production of **2** was conducted as described for **7** and **8**, but incubation was only for 36 hours. **2** was purified from the acidified culture extract by silica chromatography (toluene-acetic acid 9:10 $^{1}H$ and $^{13}C$ NMR data obtained were consistent with those determined previously for **2**.[2]

### MIC and MBC determinations

MICs were determined by broth microdilution in 96-well microtiter plates according to the CLSI guidelines.[3, 4] Cells growing in exponential phase were diluted to *ca.* $10^5$ CFU/ml into cation-adjusted Mueller-Hinton broth (MHB) before the addition of methylenomycin compounds in increasing concentrations. *Enterococci* strains were grown and diluted with Medium 92 (Trypticase soy broth 30 g, yeast extract 3 g, distilled water up to 1L). Strains were incubated for 20 h (or up to 48 h in the case of *Streptomyces* spp. and yeast) before visual inspection for growth.

Minimum bactericidal concentrations (MBCs) were determined by sub-culturing wells from MIC assays onto antibiotic-free agar plates and incubating overnight at 36 °C or at 30 °C for 4 days in the case of *Streptomyces* spp. The MBCs were further confirmed by resazurin-reduction assay with AlamarBlue™ dye as described previously.[5,6] Sequential passage of *E. faecium* 64/3 through pre-methylenomycin C lactone (**5**) and vancomycin was carried out as described by Ling *et al.*[4]

### Methods for construction of modified cosmids, plasmids, and strains

#### *Construction of plasmids pCC003 - pCC015 and introduction into S. coelicolor M145 to generate S. coelicolor W89, W95, W86, W100 and W108 respectively*

Putative biosynthetic genes were inactivated on the cosmid C73-787, which contains the entire methylenomycin biosynthetic gene cluster as well as an integrative cassette, via in-frame scar deletions and *mmyR* was replaced by an apramycin resistance cassette using PCR-targeting methodology (Figure S1).[7] The cosmid was derived from the *S. coelicolor* SCP1 ordered cosmid library.[8] The apramycin resistance cassette was amplified from pW60[9] using the forward primers A, B, C, D, and E and the reverse primers A′, B′, C′, D′ and E′ (Table S2) for *mmyR*, *mmyD*, *mmyE*, *mmyF* and *mmyO* replacement, respectively. After the separate replacement of *mmyD*, *mmyE*, *mmyF* and *mmyO* with the resistance cassette gene, the *aac(3)-IV* gene was removed using the flip recombinase,[7] leaving an in-frame scar deletion between the start and stop codon of each gene. The same methodology was used to replace the transcriptional repressor, *mmyR*, with the apramycin resistance cassette in each resulting construct. The resulting cosmids lacking *mmyR* and with deletions in *mmyD*, *mmyE*, *mmyF,* or *mmyO* were named pCC012, pCC013, pCC014, and pCC015, respectively. The construct with only *mmyR* replaced by the apramycin resistance cassette was named pCC003. The desired deletion of genes on cosmid C73-787 was confirmed by PCR using primer pairs F/ F′, G/G′, H/H′, I/I′ and J/J′ (Table S3), which are complementary to regions upstream and downstream of *mmyR*, *mmyD*, *mmyE*, *mmyF* and *mmyO*, respectively. The primers K/K′, L/L′, M/M′ and N/N′ (Table S4) were used to sub-clone *mmyD*, *mmyE*, *mmyF* and *mmyO* genes into pOSV556 to generate pCC016, pCC017, pCC018 and pCC019, respectively.[9]

pCC003 and each of pCC012-pCC015 was then introduced separately into *S. coelicolor* M145 (a derivative of *S. coelicolor* A 3(2) lacking SCP1 and SCP2) *via* conjugation with *E. coli* ET12567/pUZ8002.[7,10] This generated *S. coelicolor* W89 (*mmyR::apr*), W95 (Δ*mmyD* and *mmyR*::*apr*), W86 (Δ*mmyE* and *mmyR*::*apr*), W108 (Δ*mmyF* and *mmyR*::*apr*) and W100 (Δ*mmyO* and *mmyR*::*apr*). The engineered strains (W95, W86, W108, W100) were genetically complemented by integration of pCC016, pCC017, pCC018 and pCC019 within their genomic DNA, respectively, to generate strains W118 (W95 + *mmyD*), W115 (W86 + *mmyE*), W109 (W108 + *mmyF*) and W113 (W100 + *mmyO*) by conjugation with *E. coli* ET12567/pUZ8002 carrying each plasmid.[7,10] These strains were analysed by PCR (figure S2 and S3) and Southern blot hybridization to confirm that genetic disruption and complementation were correctly introduced. The *S. coelicolor* strains and plasmids used are summarized in Table S5.

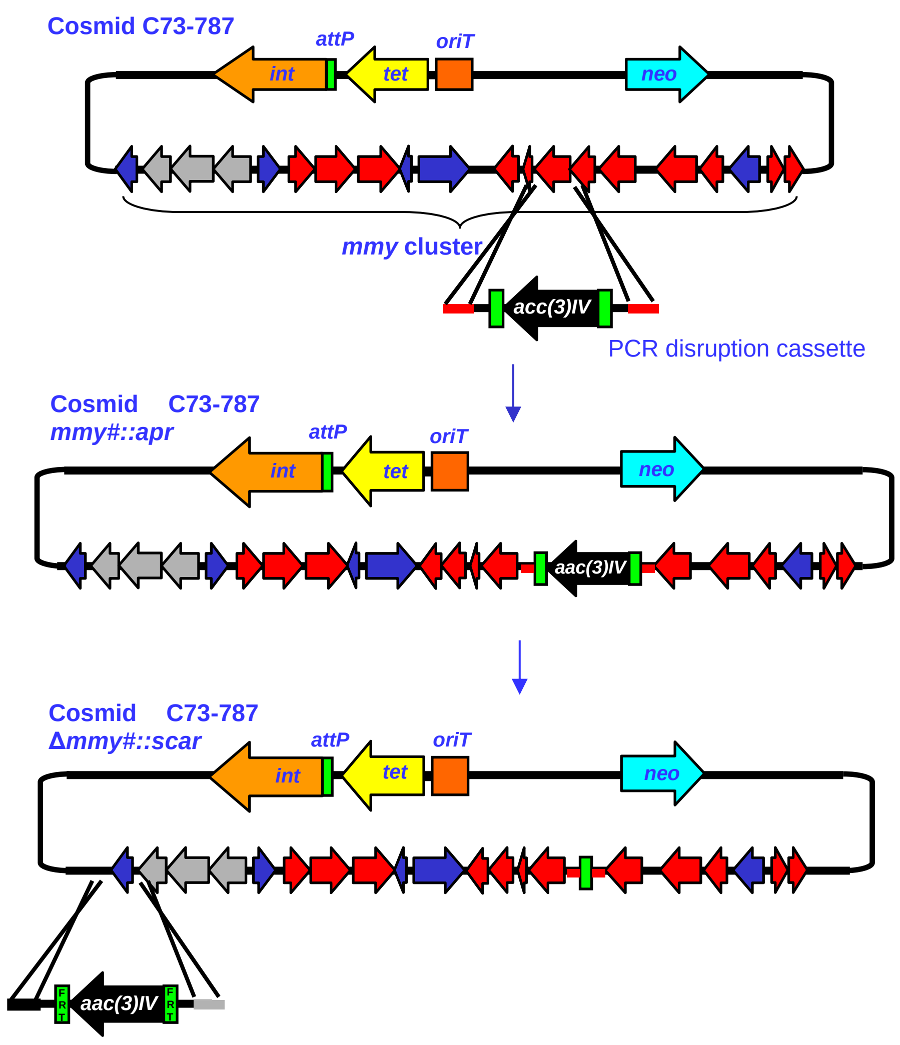


**Figure S1**. Schematic of PCR targeting method used to engineer cosmid C73-78. A PCR-generated disruption cassette containing the *aac(3)-IV* gene, which confers apramycin resistance, flanked by flip recombinase target (FRT) sites is inserted between the start and stop codon on the the gene of interest (*mmy#*). The flip recombinase is then used to excise the *aac(3)-IV* gene, leaving an 81 bp in frame “scar” sequence between the start and stop codons. Finally, the *mmyR* gene is replaced with a PCR-generated disruption cassette containing the *aac(3)-IV* gene.

**Table S1**. Putative functions of proteins encoded by the methylenomycin biosynthetic gene cluster

| **Protein** | **Homologue (% identity)** | **Proposed function** |
|---|---|---|
| MmyA | RedQ *Streptomyces coelicolor* (33) | Acyl carrier protein |
| MmyC | FabH *Escherichia coli* (39) | β-ketoacyl synthase III |
| MmyD | AvrD *Pseudomonas syringae* (28) | Butenolide synthase |
| MmyE | PlmM *Streptomyces* sp. HK803 (32) | Flavin-dependent enoyl reductase |
| MmyF | NtaB *Chelatobacter heintzii* (40) | Flavin reductase, partner to monooxygenase |
| MmyG | JadP *Streptomyces venezuelae* (29) | NAD(P)-dependent dehydrogenase |
| MmyK | Adk *Escherichia coli* (30) | Kinase |
| MmyO | LimB *Rhodococcus erythropolis* (43) | Flavin-dependent monooxygenase |
| MmyP | SsgB *Streptomyces griseus* (30) | Phosphatase |
| MmyQ | NdpG *Rhodococcus opacus* (30) | Coenzyme F-420-dependent reductase |
| MmyT | MtmZ *Streptomyces argillaceus* (31) | Thioesterase |
| MmyX | MmyK *Streptomyces coelicolor* (42) | Kinase |
| MmyY | JadX *Streptomyces venezuelae* (27) | Unknown – belongs to NTF2-like superfamily |

**Table S2**. PCR primers used to create disruption cassettes for *mmyD*, *mmyE*, *mmyF*, *mmyO*, and *mmyR*

| **Primers** | **Sequence** |
|---|---|
| A | 5’_CCCGTTTTCTCACGACCTTGAGAGGACTCGGGCGTTGTGTATTCCGGGGATCCGTCGACC_3’ |
| A′ | 5'_CCGGAGCTCGTTGTCCGCCGGATCCTGGTCGCCGCAGTCATGTAGGCTGGAGCTGCTTC_3' |
| B | 5’_ATCGATTGCACCTGTCGGGAAAAAACTGGAGGGTGCATGATTCCGGGGATCCGTCGACC_3’ |
| B′ | 5'_CGTGTCCTGCAGGGCAGGCCGACGGTGGACAGTGGGTCATGTAGGCTGGAGCTGCTTC_3' |
| C | 5’_GGCCCCCCACCGGGAACCAGTCATCCGAAGGGACAGATGATTCCGGGGATCCGTCGACC_3’ |
| C′ | 5'_CGGACCCGGGCCGTGGTGTCACCGCCCTGCACGGCGTCATGTAGGCTGGAGCTGCTTC_3' |
| D | 5’_GGCTGACTGTTTCCCCTTCTCCTCCAGGGAGTCCGCATGATTCCGGGGATCCGTCGACC_3’ |
| D′ | 5'_CCTGAGCTCCATGGCGGACGAACTGCCGTCAGGTCCTCATGTAGGCTGGAGCTGCTTC_3' |
| E | 5’_CCCACGCTGCAATTTCAGCGCGACCTTGAGCTGATAGAAATTCCGGGGATCCGTCGACC_3’ |
| E′ | 5'_TCGCCGGGTGGAGCCGGTGAAGTGCGGGGCGACGTAGCGTGTAGGCTGGAGCTGCTTC_3' |

Underlined sequences correspond to the 20 nucleotides P1 and 19 nucleotides P2 sequences described in the procedure developed by Gust *et al*.[7]

**Table S3**. PCR primers used to screen transconjugants for integration of each engineered cosmid into the chromosome of *S. coelicolor* M145

| Primers | Sequence |
|---|---|
| F | 5’_GCCATCGGTTGAATCCTG_3’ |
| F′ | 5'_CAGGAAACGGACTGCCTG_3' |
| G | 5’_CCGGCAATGACGAAATAG_3’ |
| G′ | 5’_TGGCCAGGTTCATAGGAG_3’ |
| H | 5’_CACCGGGAACCAGTCATC_3’ |
| H′ | 5'_GCTTGCCTCACCGAGTTG_3' |
| I | 5'_CCCGCTGCCATGCGATTC_3' |
| I′ | 5’_ACCGCTACGGCCTAGTGC_3’ |
| J | 5’_GTGCACGGTTTACGGGATGAG_3’ |
| J′ | 5’_ACGCCGATGCGTATCGGTTC_3’ |

**Table S4**. PCR primers used to amplify *mmyD*, *mmyE, mmyF* and *mmyO* for cloning into pOSV556

| Primers | Sequence |
|---|---|
| K | 5’_GGGGGAAGCTTGAGAAGGGAGCGGACAT**ATG**CCAGTCAGCGGTTCCC_3’ |
| K′ | 5'_GATAATCTCGAGGTGGACAGTGGGTCAAG_3' |
| L | 5’_AATGCTCACGATTGTGAGGAGGGGCAG**ATG**CACG_3’ |
| L′ | 5’_AATAATAAGCTGCCCTGCACGGCGTCAG_3’ |
| M | 5’_AAAGGAAGCTTAGGAGGGTCCGC**ATG**GCTACG_3’ |
| M′ | 5'_GGGAAACTCGAGCCGTCAGGTCCTCATG_3' |
| N | 5'_AAAGGGAAGCTTAGGAGGACGTTC**ATG**TACCCCG_3' |
| N′ | 5’_GGGAAACTCGAGGGCGTGCACGGTTTAC_3’ |
| O | 5’_ATATAGGATCCAGGAGGAACAGCATGACCACTG_3’ |
| O′ | 5’_ ATTATTAAGCTTGGCAGTGTCCAGGAGCG_3′ |

Underlined sequences correspond to restriction sites (K, L’, M, N – *Hin*dIII; K’, M’, N’ – *Xho*I; L – *Ale*I)

**Table S5**. List of plasmids and strains

| Strains / Plasmids | Relevant properties | Reference |
|---|---|---|
| *S. coelicolor* strains | | |
| M145 | SCP1$^-$, SCP2$^-$ (methylenomycin non-producer) | 10 |
| W89 | M145 containing pCC003::*attB* → *mmyR*::*apr* | This study |
| W95 | M145 containing pCC012::*attB* → *mmyR*::*apr* and Δ*mmyD* | This study |
| W86 | M145 containing pCC013::*attB* → *mmyR*::*apr* and Δ*mmyE* | This study |
| W108 | M145 containing pCC014::*attB*→ *mmyR*::*apr* and Δ*mmyF* | This study |
| W100 | M145 containing pCC015::*attB*→ *mmyR*::*apr* and Δ*mmyO* | This study |
| W118 | W95 containing pCC016::*attB′*→ *mmyD* complementation | This study |
| W115 | W86 containing pCC017::*attB′*→ *mmyE* complementation | This study |
| W109 | W108 containing pCC018::*attB′*→ *mmyF* complementation | This study |
| W113 | W100 containing pCC019::*attB′*→ *mmyO* complementation | This study |
| W110 | M145 containing pOSV556/*mmyOF* integrated into *attB′* | This study |
| W301 | W110 containg pIJ86/*mmr* | This study |
| W302 | M145 with pIJ86/*mmr* | This study |
| Plasmids | | |
| C73_787 | Integrative cosmid containing the *mmy* gene cluster | 8 |
| pCC003 | C73_787 with *mmyR*::*apr* | This study |
| pCC004 | C73_787 with *mmyD::apr* | This study |
| pCC005 | C73_787 with *mmyE::apr* | This study |
| pCC006 | C73_787 with *mmyF*::*apr* | This study |
| pCC007 | C73_787 with *mmyO::apr* | This study |
| pCC008 | C73_787 with Δ*mmyD* | This study |
| pCC009 | C73_787 with Δ*mmyE* | This study |
| pCC010 | C73_787 with Δ*mmyF* | This study |
| pCC011 | C73_787 with Δ*mmyO* | This study |
| pCC012 | C73_787 with Δ*mmyD* and *mmyR*::*apr* | This study |
| pCC013 | C73_787 with Δ*mmyE* and *mmyR*::*apr* | This study |
| pCC014 | C73_787 with Δ*mmyF* and *mmyR*::*apr* | This study |

| | | |
|---|---|---|
| pCC015 | C73_787 with Δ*mmyO* and *mmyR*::*apr* | This study |
| pCC016 | pOSV556t containing *mmyD* gene and artificial RBS | This study |
| pCC017 | pOSV556t containing *mmyE* gene and artificial RBS | This study |
| pCC018 | pOSV556t containing *mmyF* gene and artificial RBS | This study |
| pCC019 | pOSV556t containing *mmyO* gene and artificial RBS | This study |
| pOSV556/ *mmyOF* | pOSV556 containing *mmyOF* under the control of the *ermE** promoter | This study |
| pIJ86/*mmr* | pIJ86 containing *mmr* under the control of the *ermE** promoter | This study |

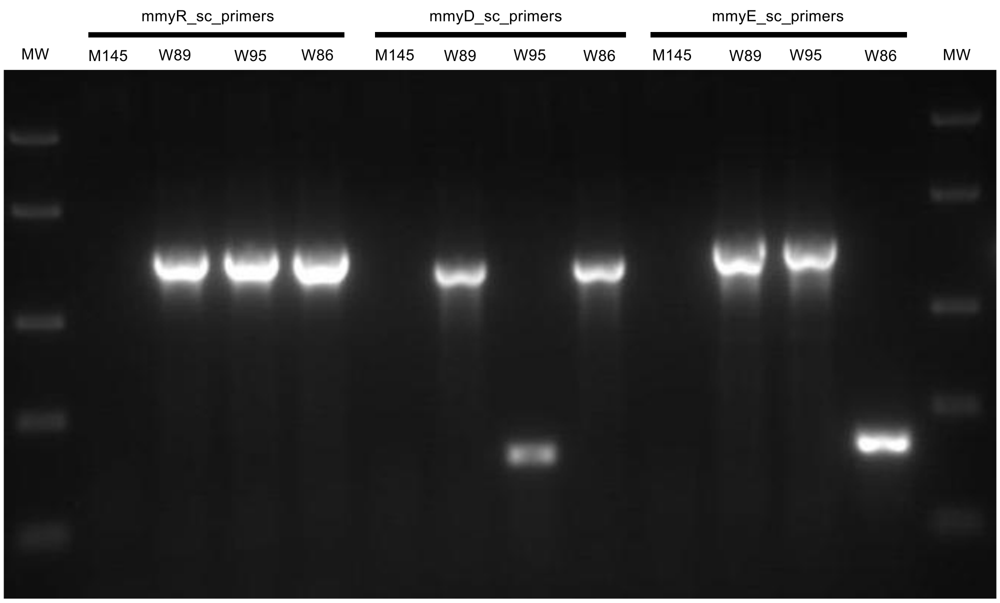


**Figure S2**. PCR analysis of genomic DNA from *S. coelicolor* M145, W89, W95 and W86.

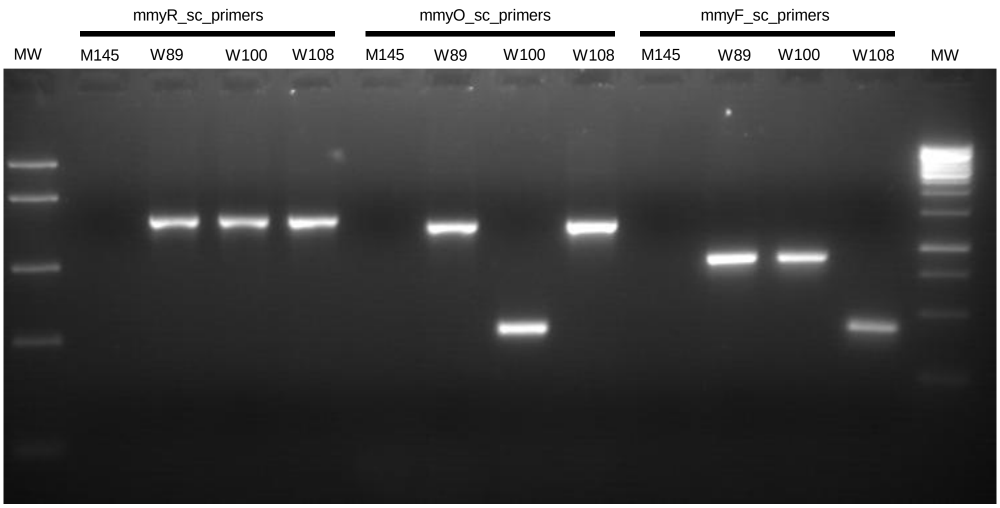


**Figure S3**. PCR analysis of genomic DNA from *S. coelicolor* M145, W89, W100 and W108

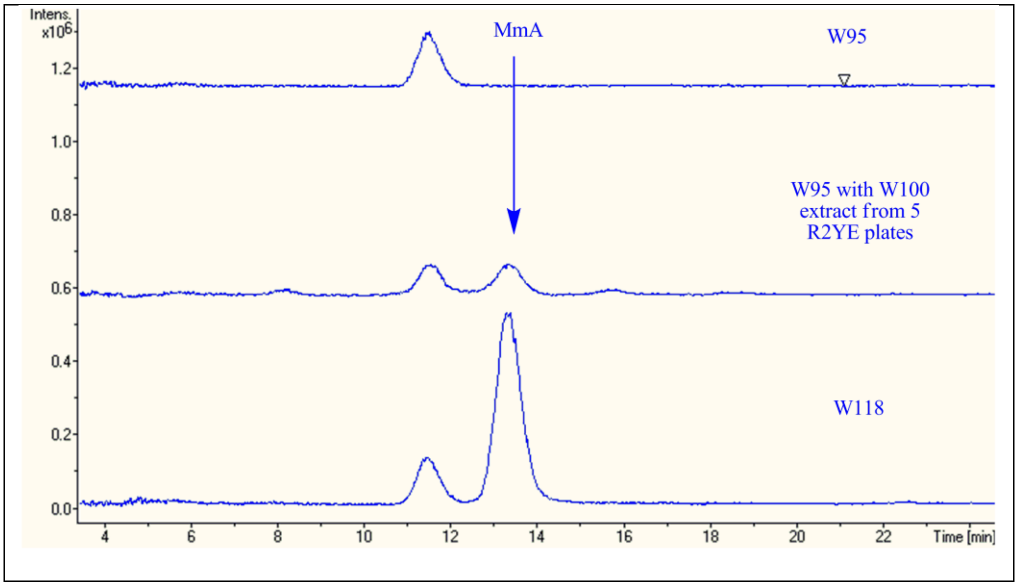


**Figure S4**. Genetic and chemical complementation of *S. coelicolor* W95 (Δ*mmyD*/*mmyR*::*apr*) restores methylenomycin A production. Extracted ion chromatogram at *m/z* = 183.0650 (corresponding to $[M+H]^+$ for **1**) from LC-MS analysis of extracts *S. coelicolor* W95 (top), *S. coelicolor* W95 fed with an organic extract of *S. coelicolor* W100 (middle), and *S. coelicolor* W118 (bottom).

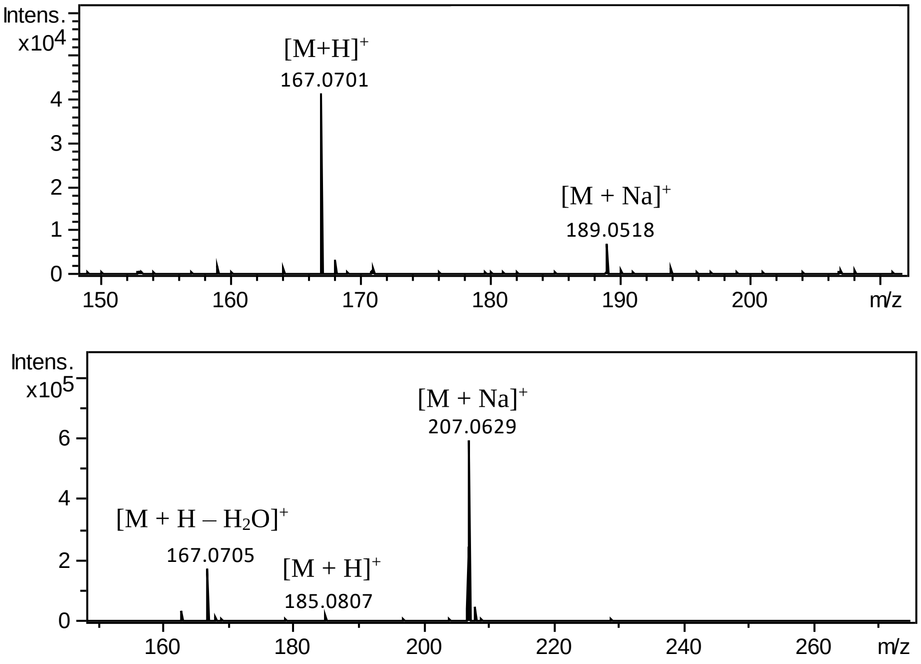


**Figure S5.** High-resolution mass spectra of pre-methylenomycin C lactone (**5**) (top) and pre-methylenomycin C (**6**) (bottom).

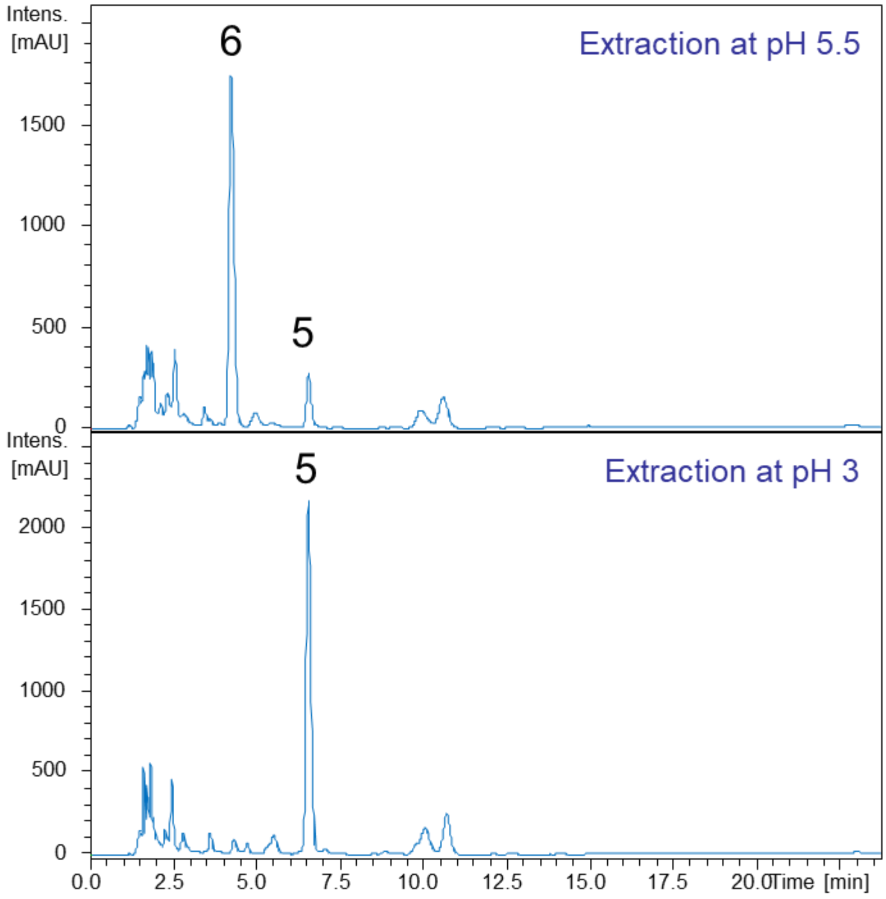


**Figure S6** UV chromatograms at 230nm from HPLC purification of neutral (pH = 5.5; top) or acidic (pH = 3; bottom) extracts of *S. coelicolor* W86.

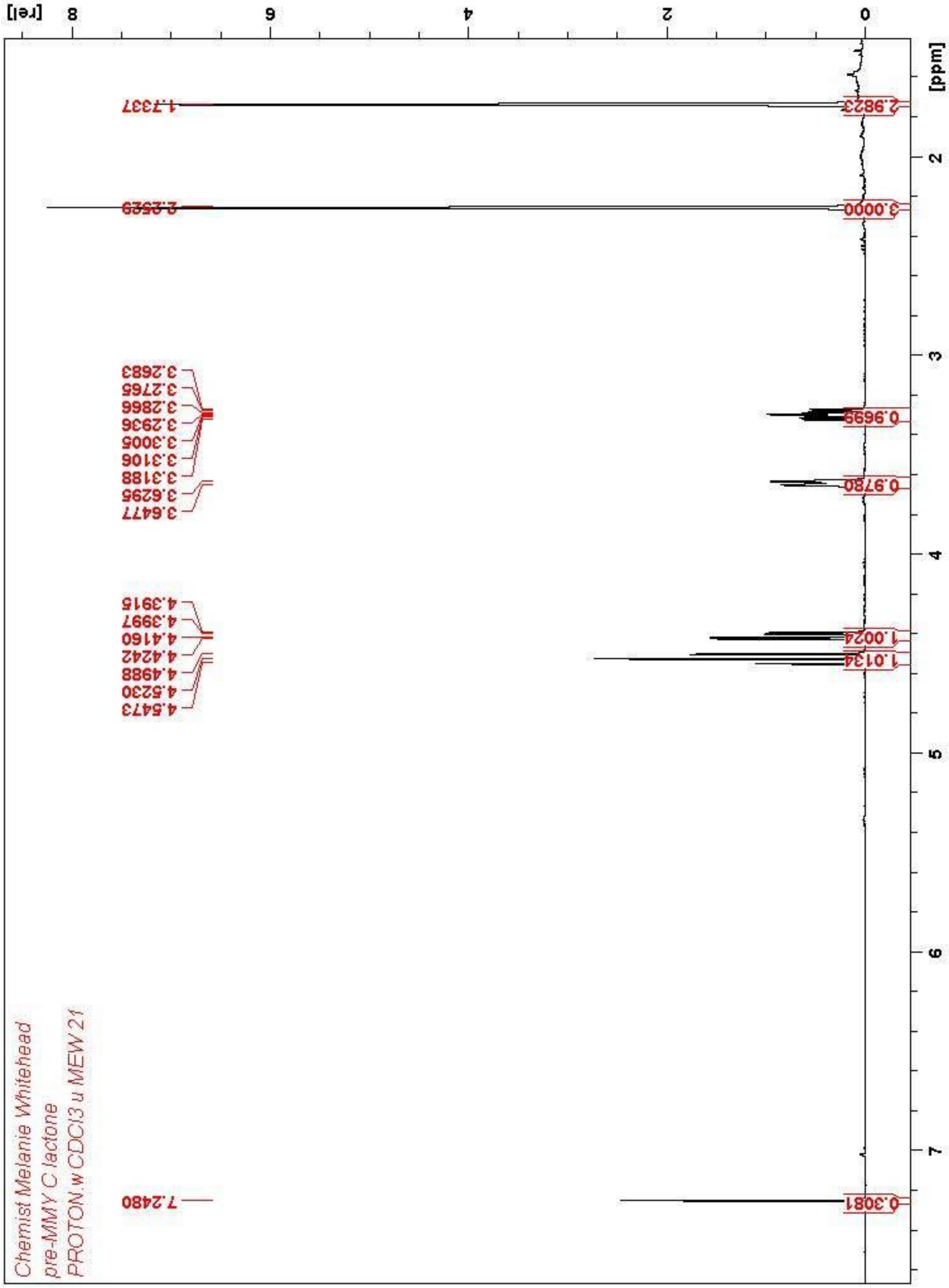


**Figure S7**. [1]H-NMR spectrum of pre-methylenomycin C lactone (**5**) in $CDCl_3$

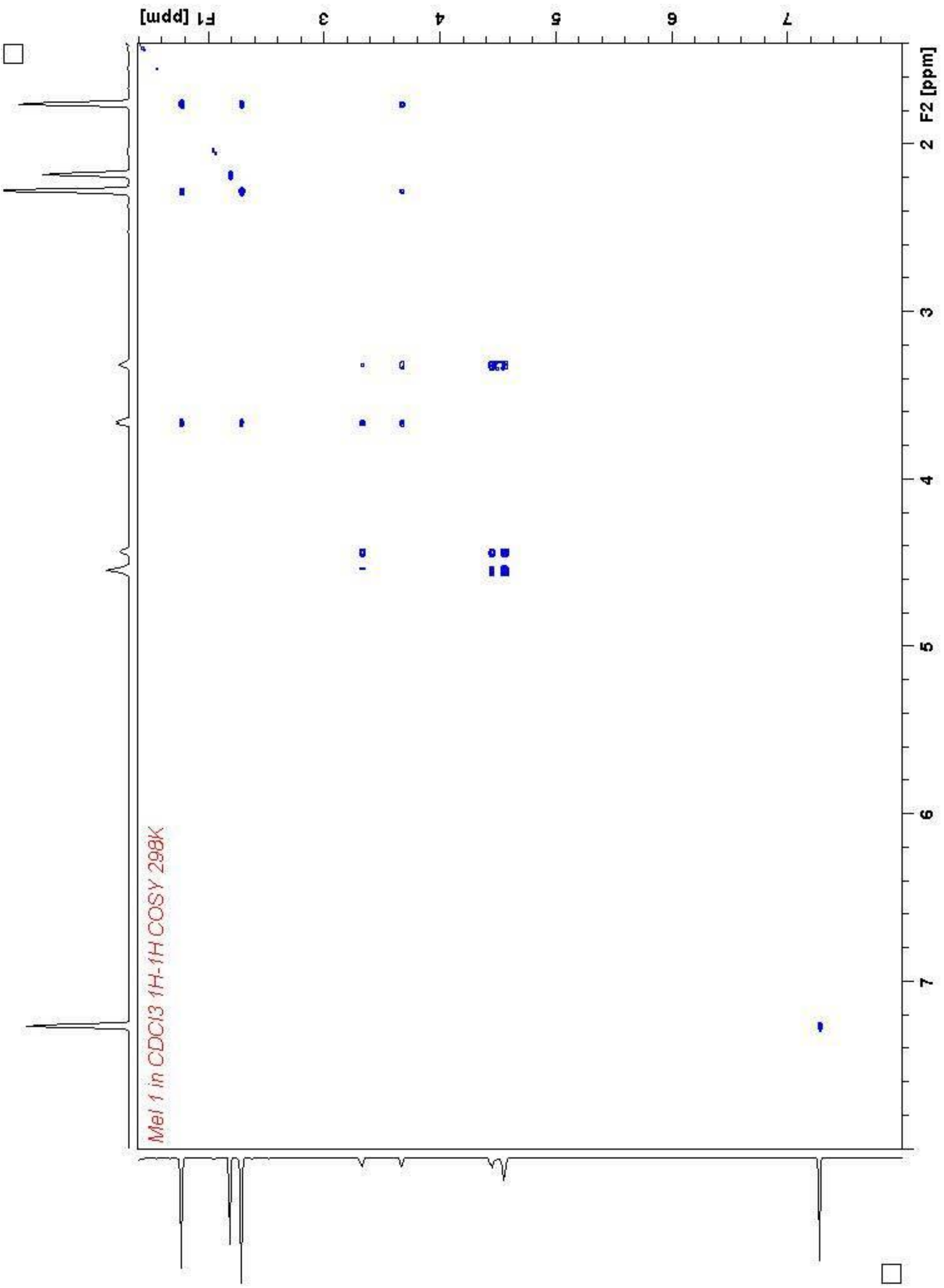


**Figure S8**. COSY spectrum of pre-methylenomycin C lactone (**5**) in $CDCl_3$

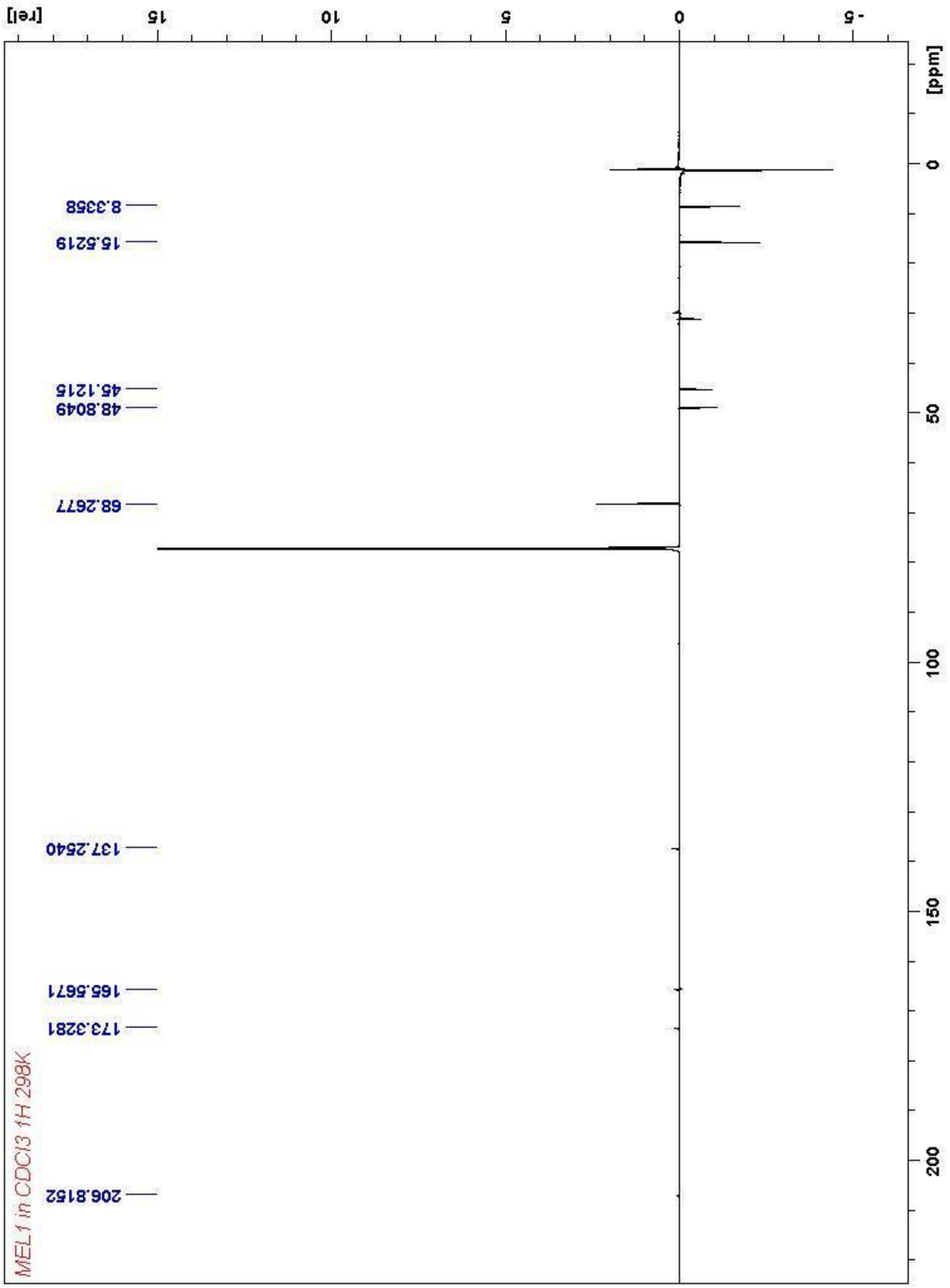


**Figure S9**. $^{13}$C-NMR spectrum of pre-methylenomycin C lactone (**5**) in $CDCl_3$

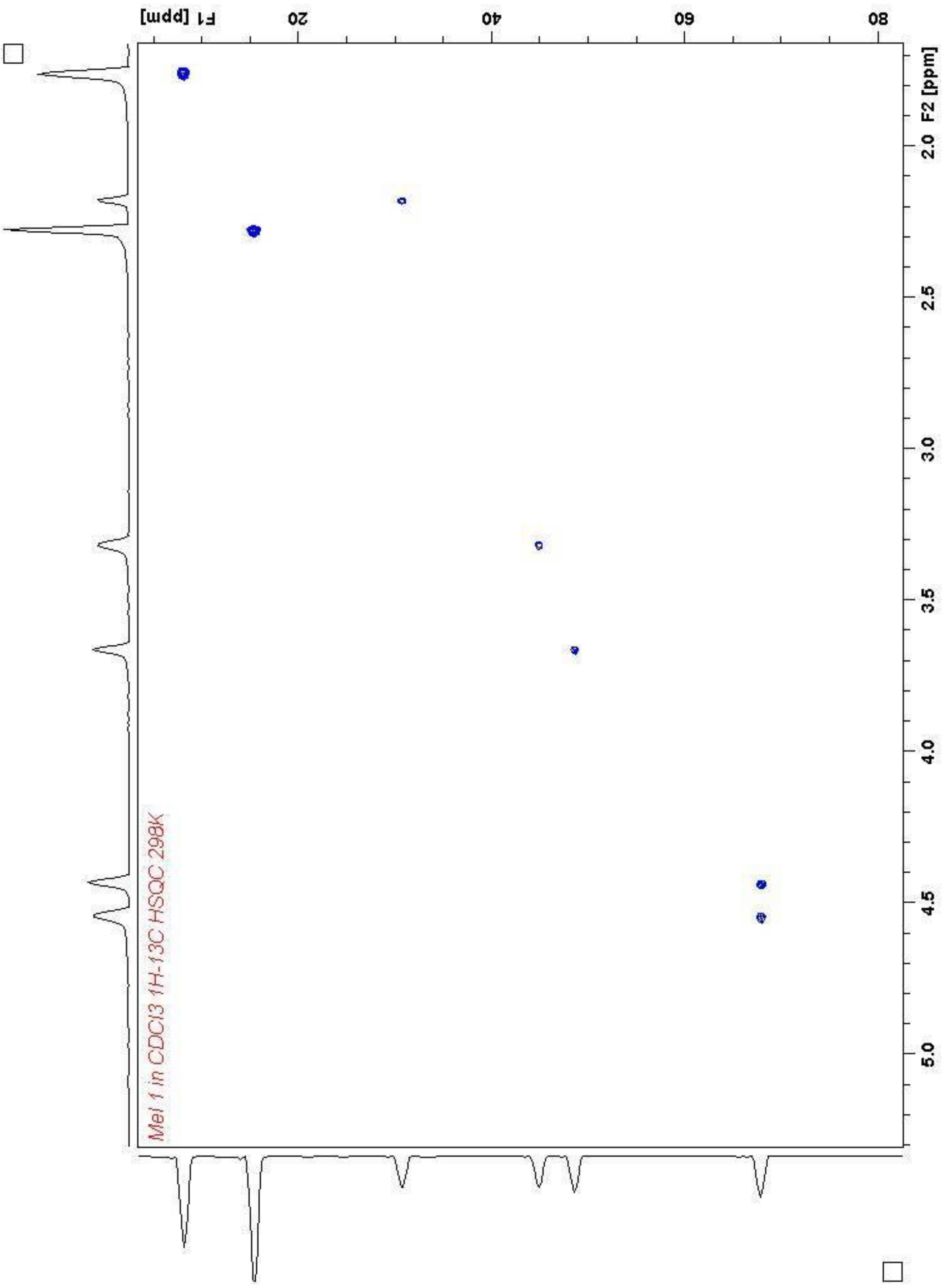


**Figure S10**. HSQC spectrum of pre-methylenomycin C lactone (**5**) in $CDCl_3$

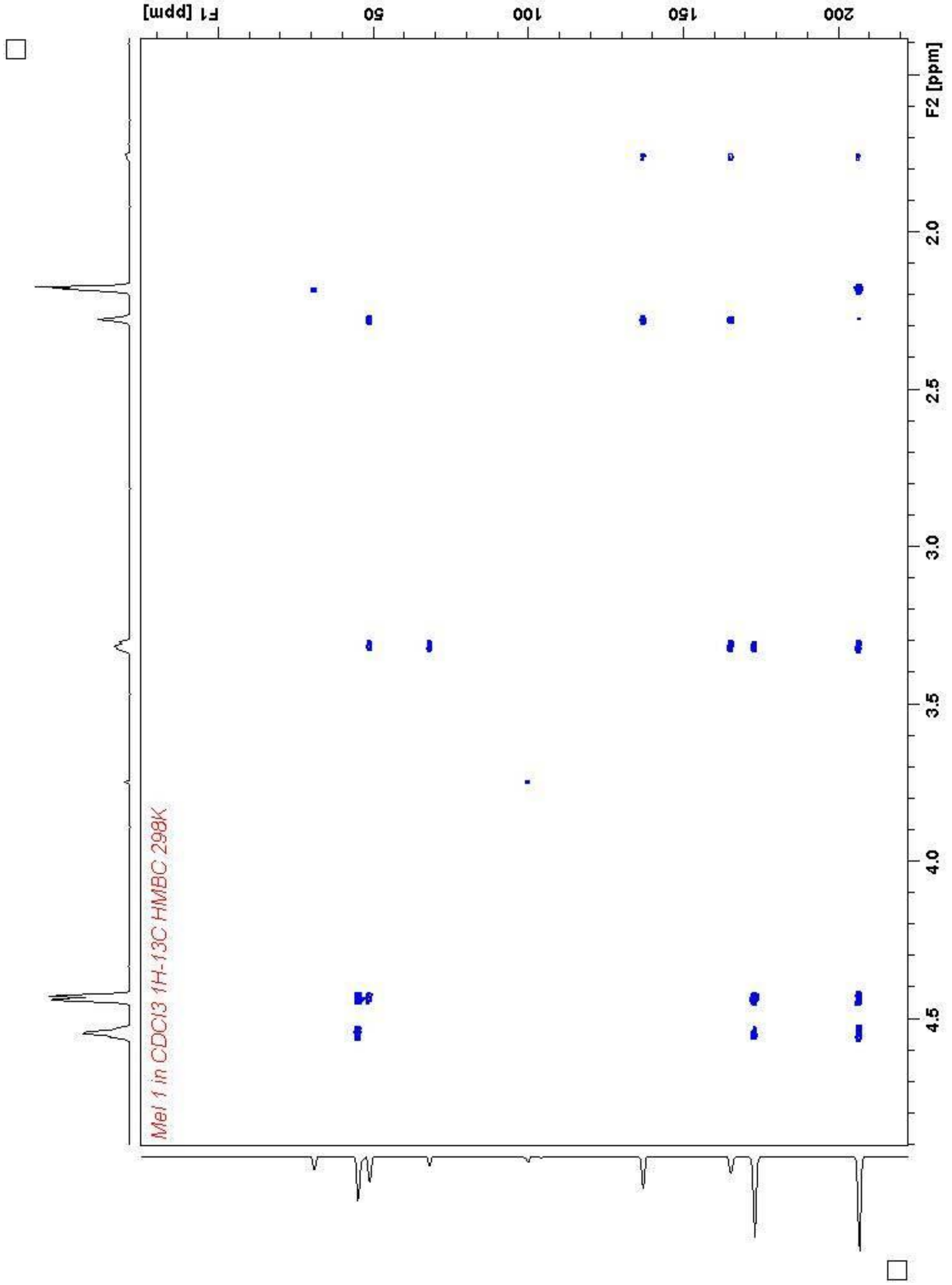


**Figure S11**. HMBC spectrum of pre-methylenomycin C lactone (**5**) in $CDCl_3$

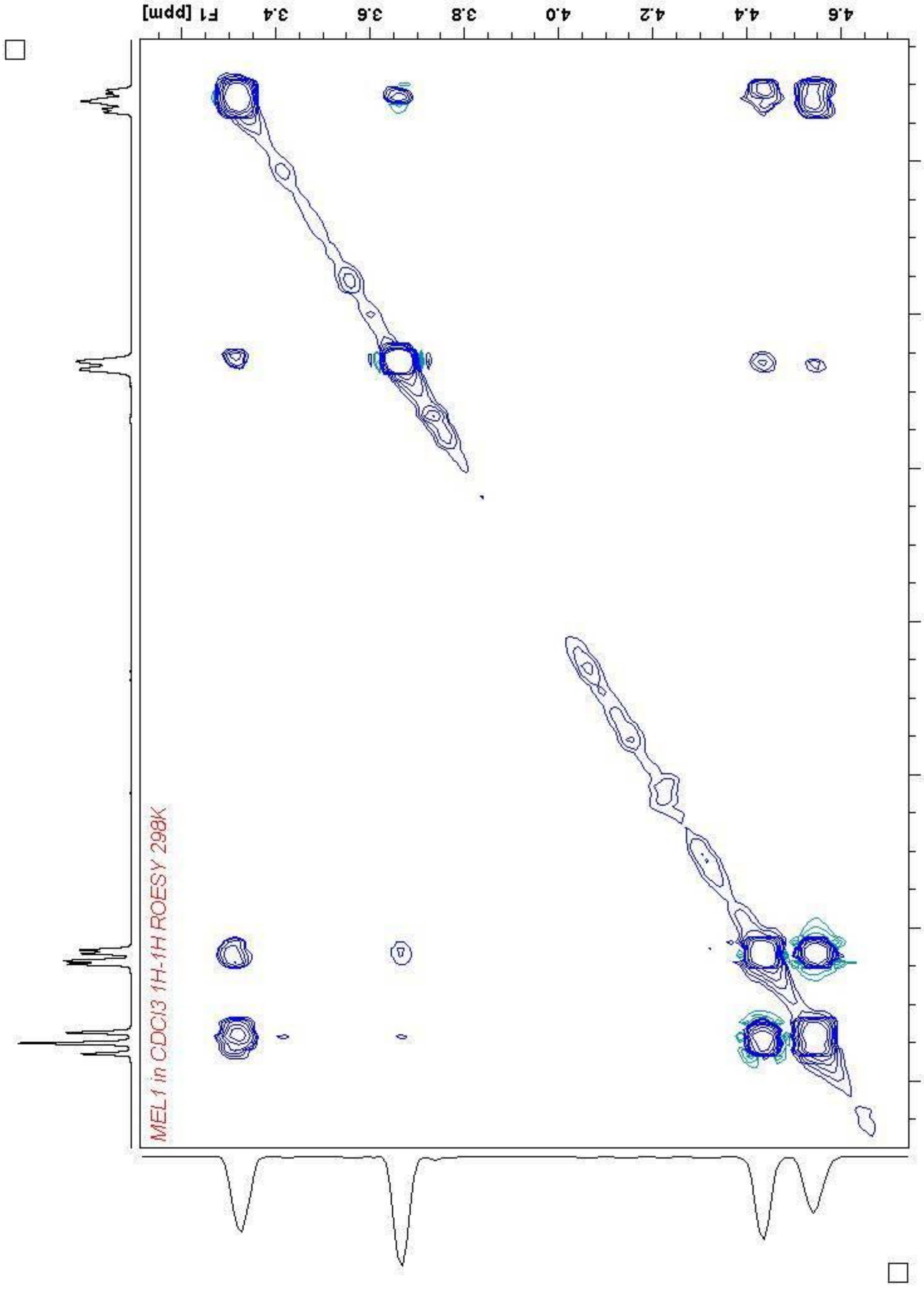


**Figure S12**. ROESY spectrum of pre-methylenomycin C lactone (**5**) in $CDCl_3$

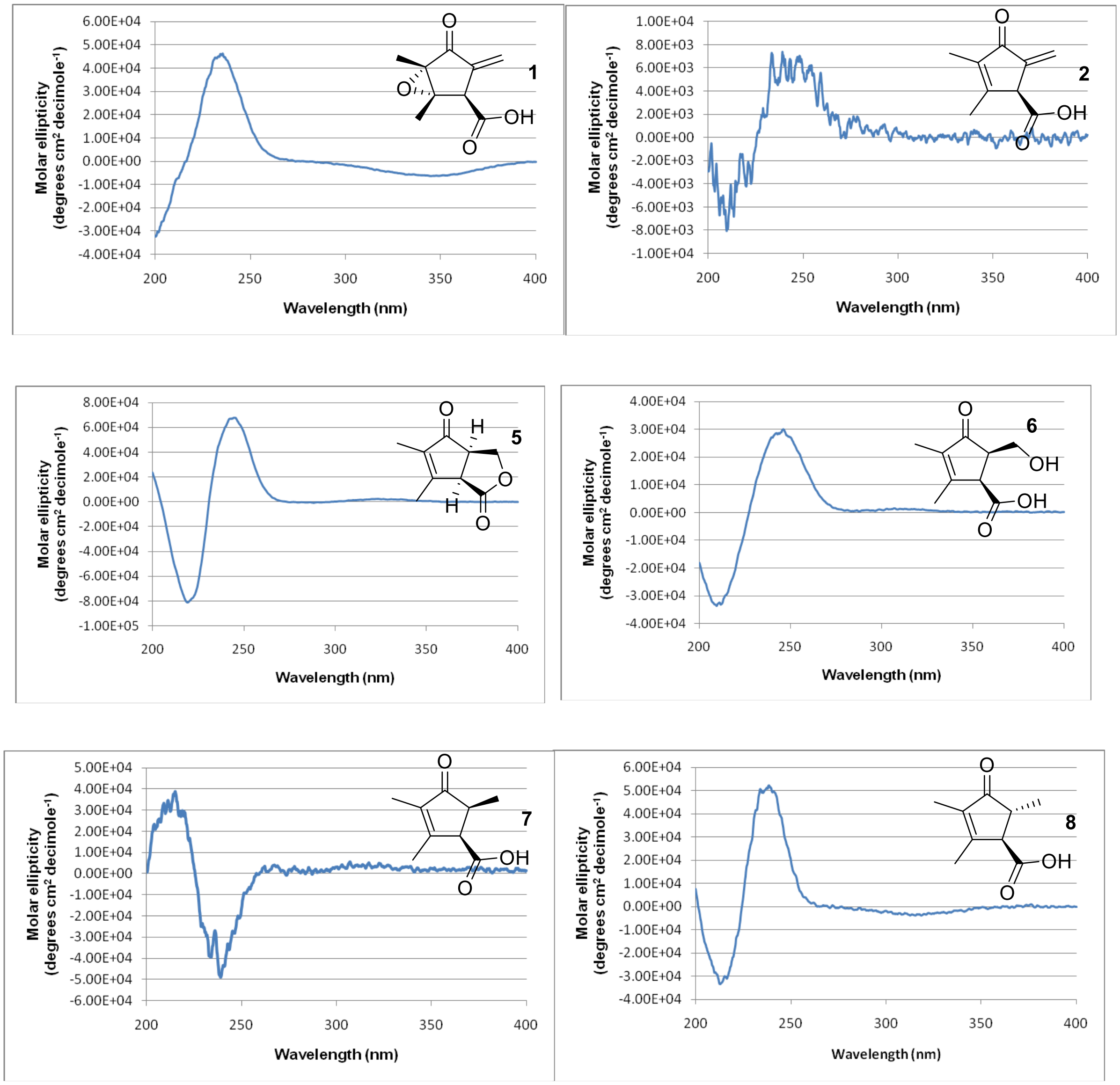


**Figure S13.** CD spectrum of methylenomycin A (**1**), methylenomycin C (**2**), pre-methylenomycin C lactone (**5**), pre-methylenomycin C (**6**), methylenomycin D1 (**7**) and methylenomycin D2 (**8**)

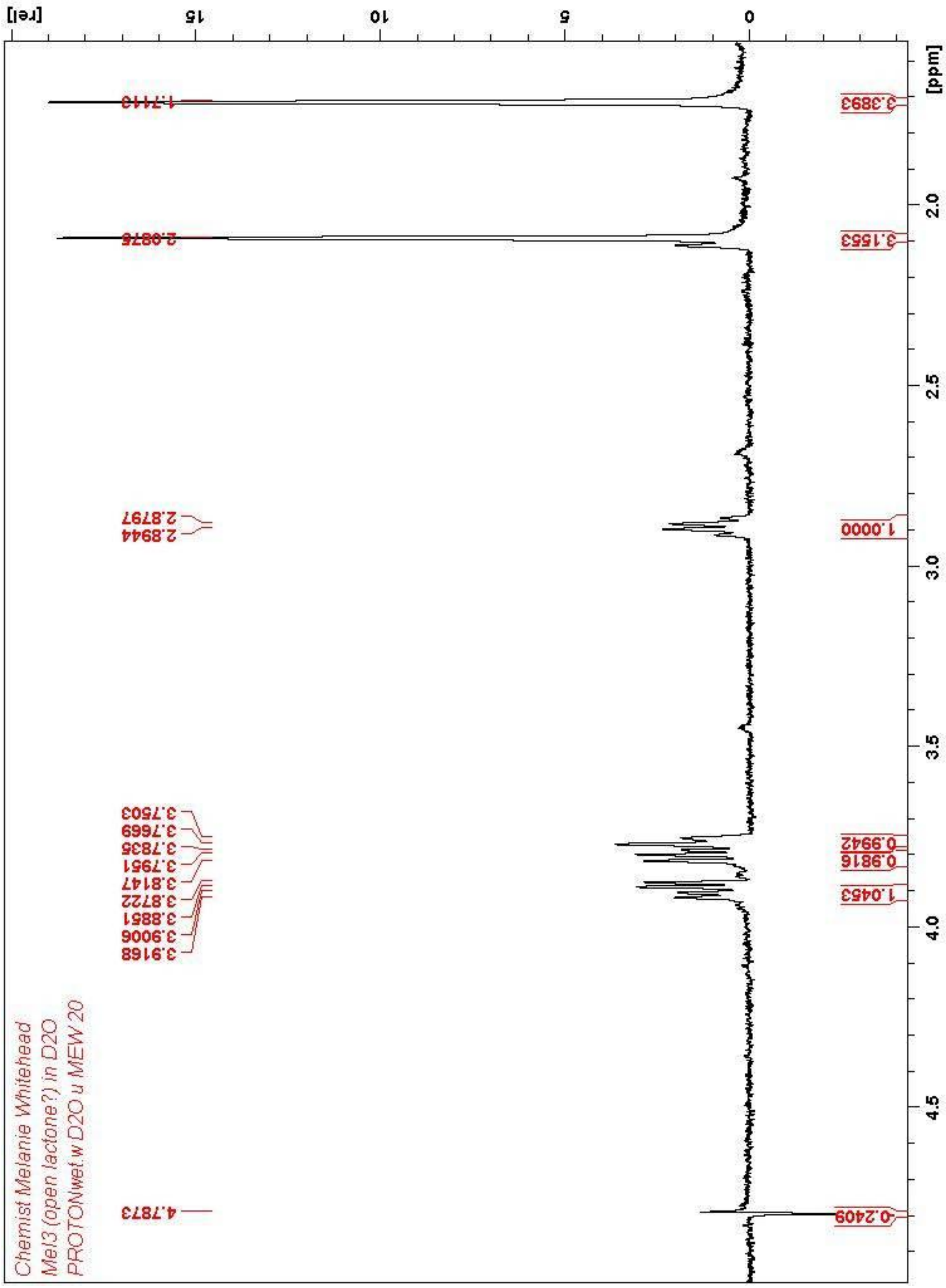


**Figure S14**. $^1$H-NMR spectrum of pre-methylenomycin C (**6**) in $D_2O$

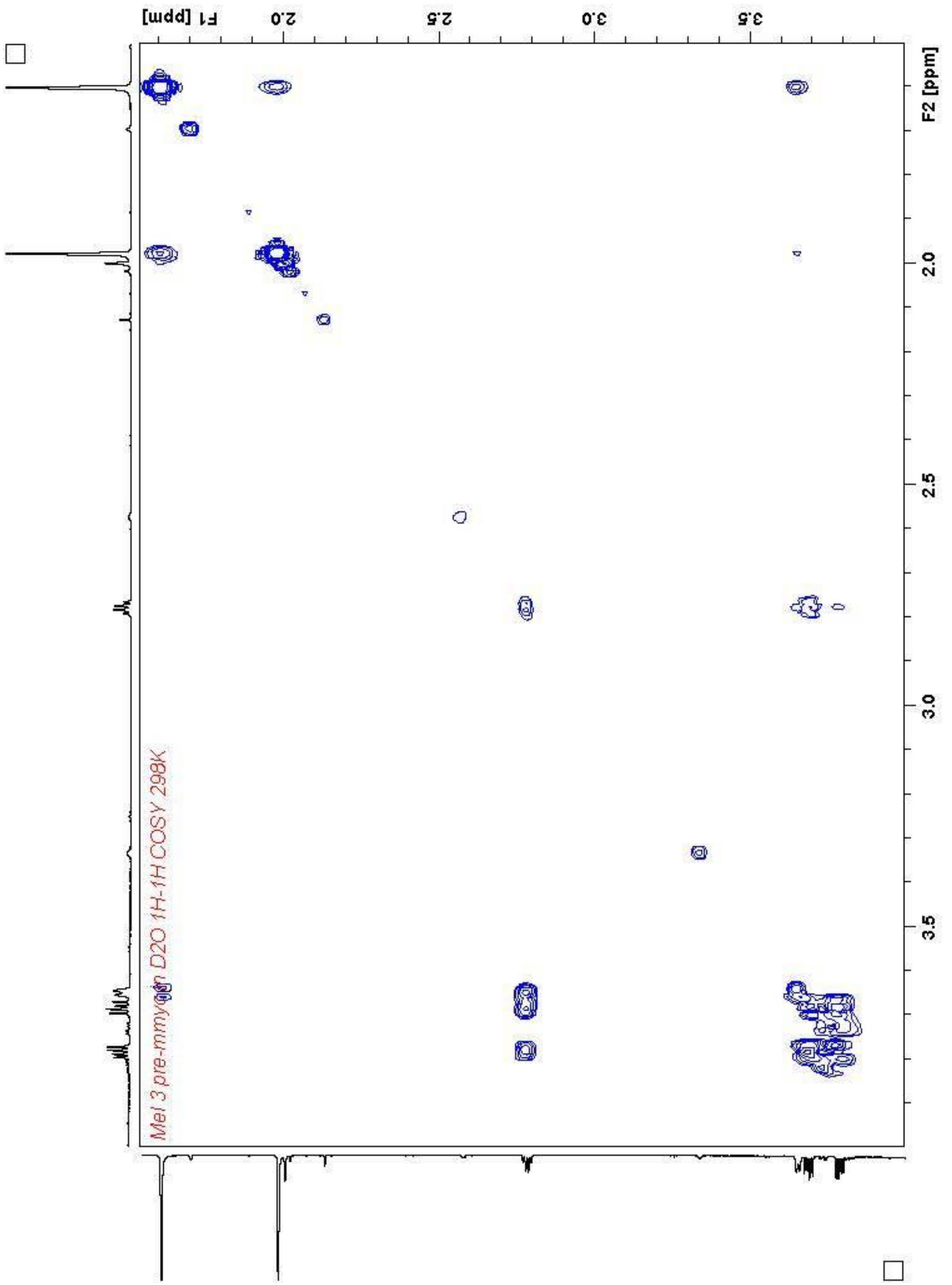


**Figure S15**. COSY spectrum of pre-methylenomycin C (**6**) in $D_2O$

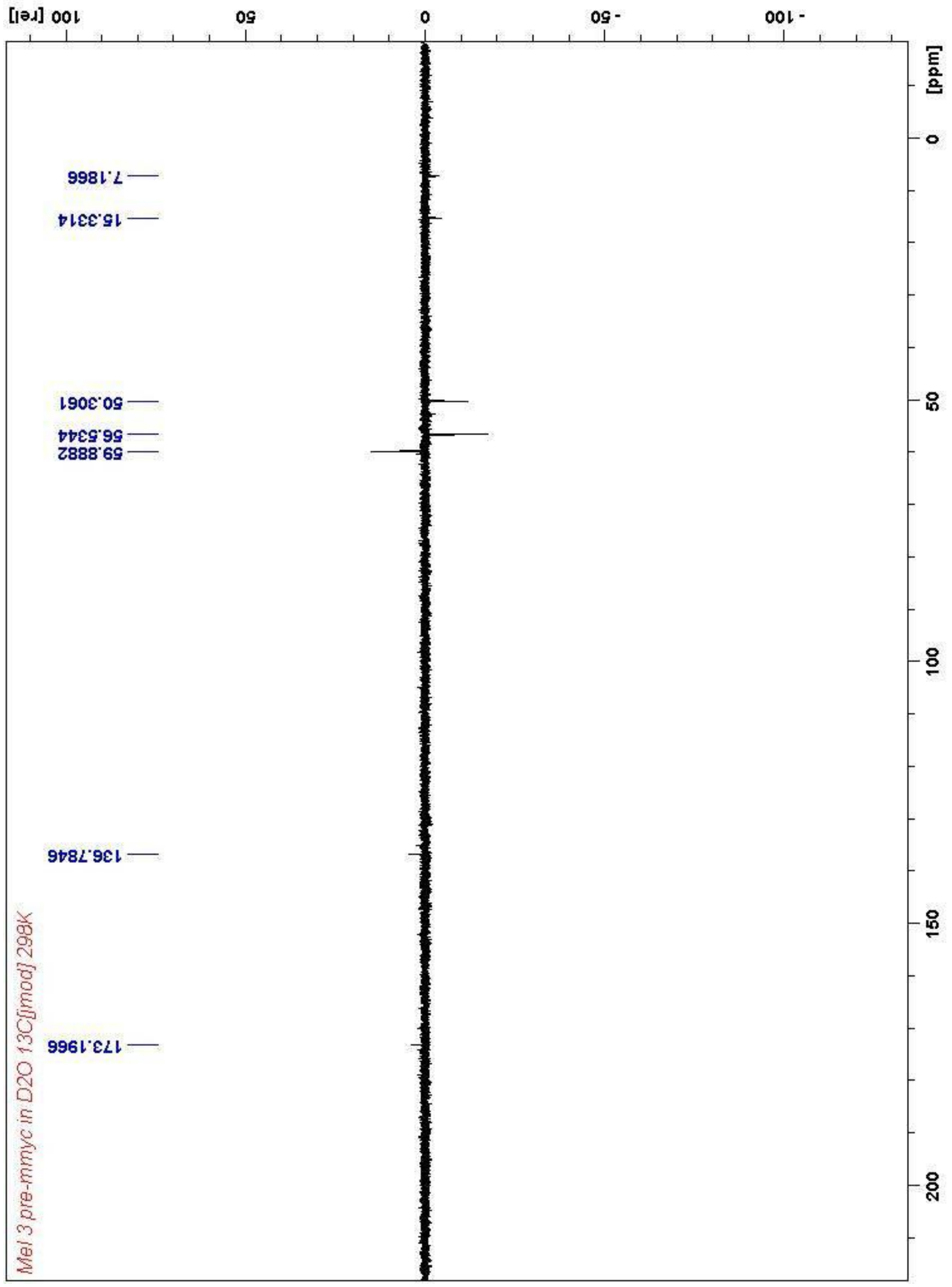

**Figure S16**. $^{13}$C-NMR spectrum of pre-methylenomycin C (**6**) in $D_2O$

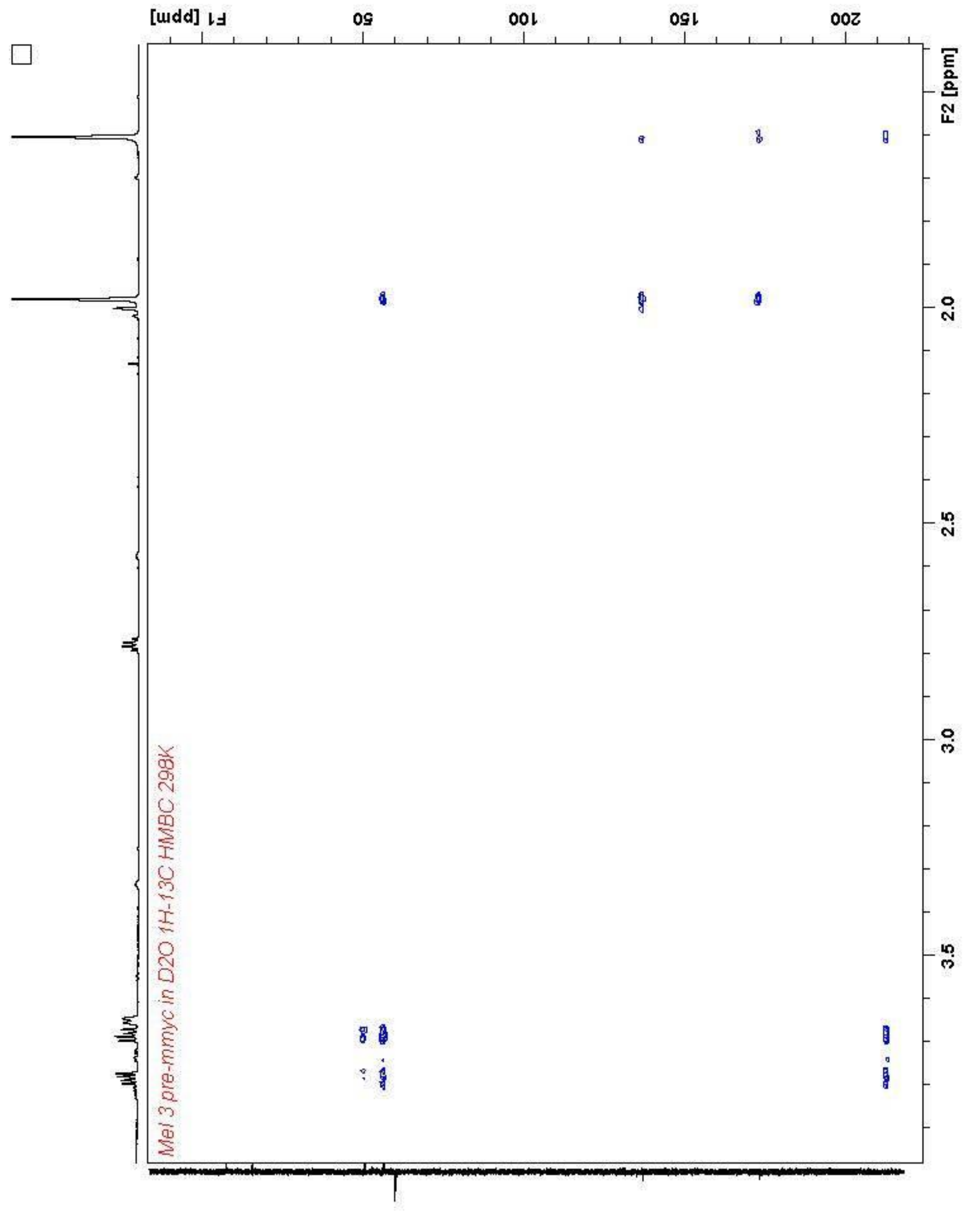


**Figure S17**. HMBC spectrum of pre-methylenomycin C (**6**) in $D_2O$

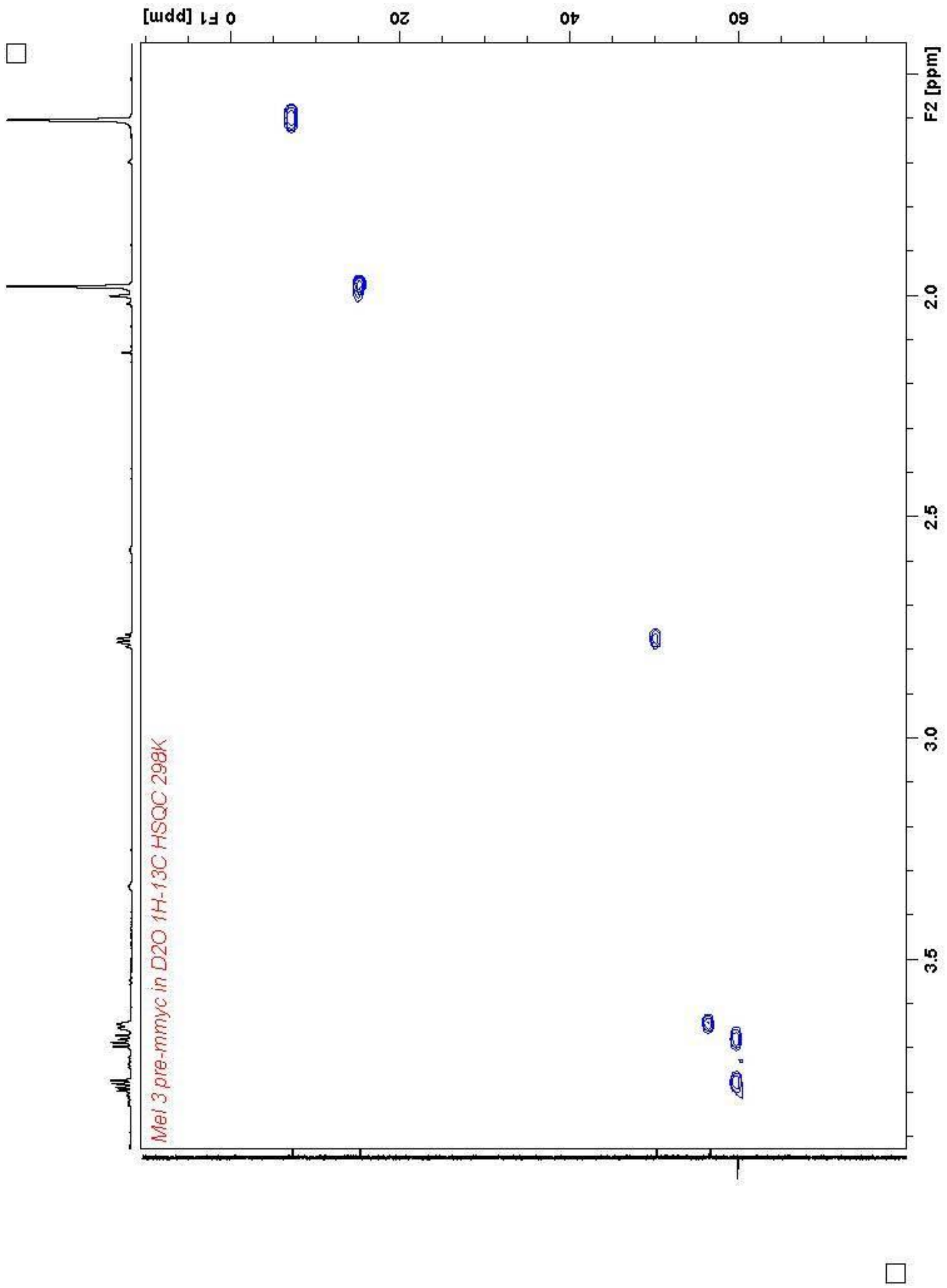


**Figure S18**. HSQC spectrum of pre-methylenomycin C (**6**) in $D_2O$

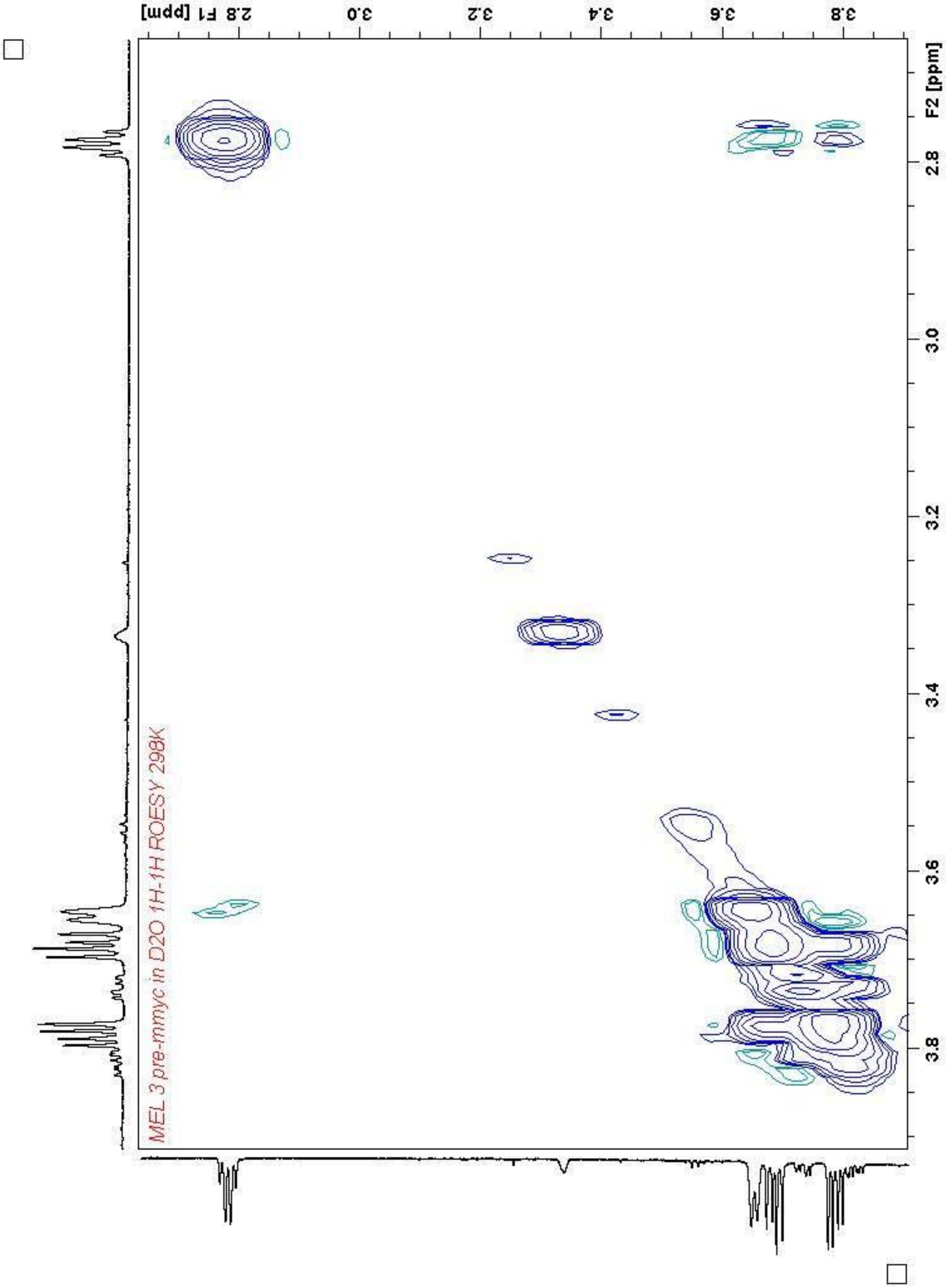


**Figure S19**. ROESY spectrum of pre-methylenomycin C (**6**) in $D_2O$

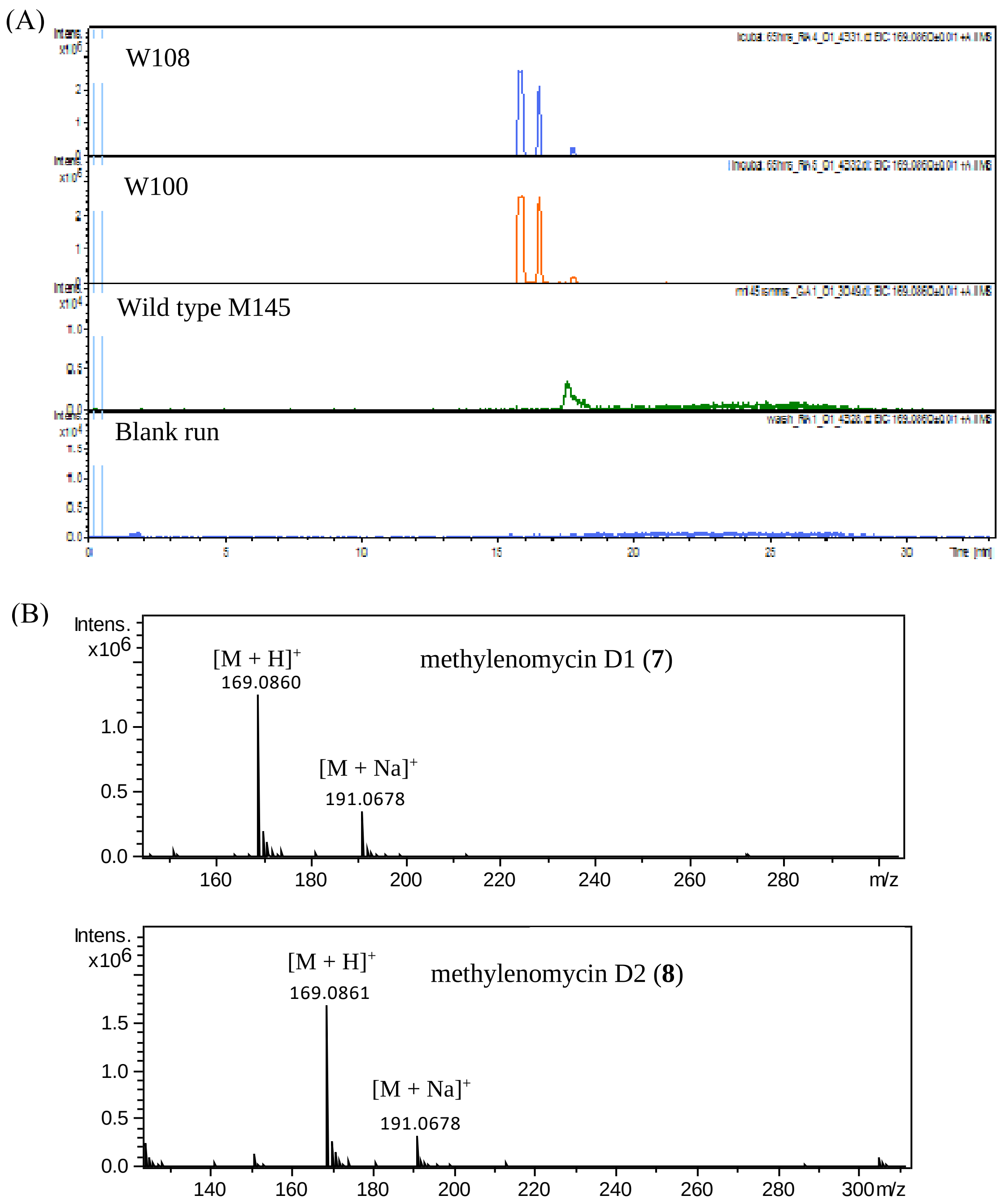


**Figure S20.** (A) Extracted ion chromatograms (EICs) at m/z = 169.0860, corresponding to [M + H$^+$] for methylenomycin D1,**7** (15.8 min) and methylenomycin D2, **8** (16.6 min), following LC-MS analyses of extracts of strain W108 (M145/C73_787/Δ*mmyF*/*mmyR*::*apr*) and W100 (M145/C73_787/Δ*mmyO*/*mmyR*::*apr*) grown for 3 days (B) High-resolution mass-spectrometry data showing m/z = 169.086 and 191.0678, corresponding to [M + H$^+$] and [M + Na$^+$] for **7** (top spectra) and **8** (bottom spectra).

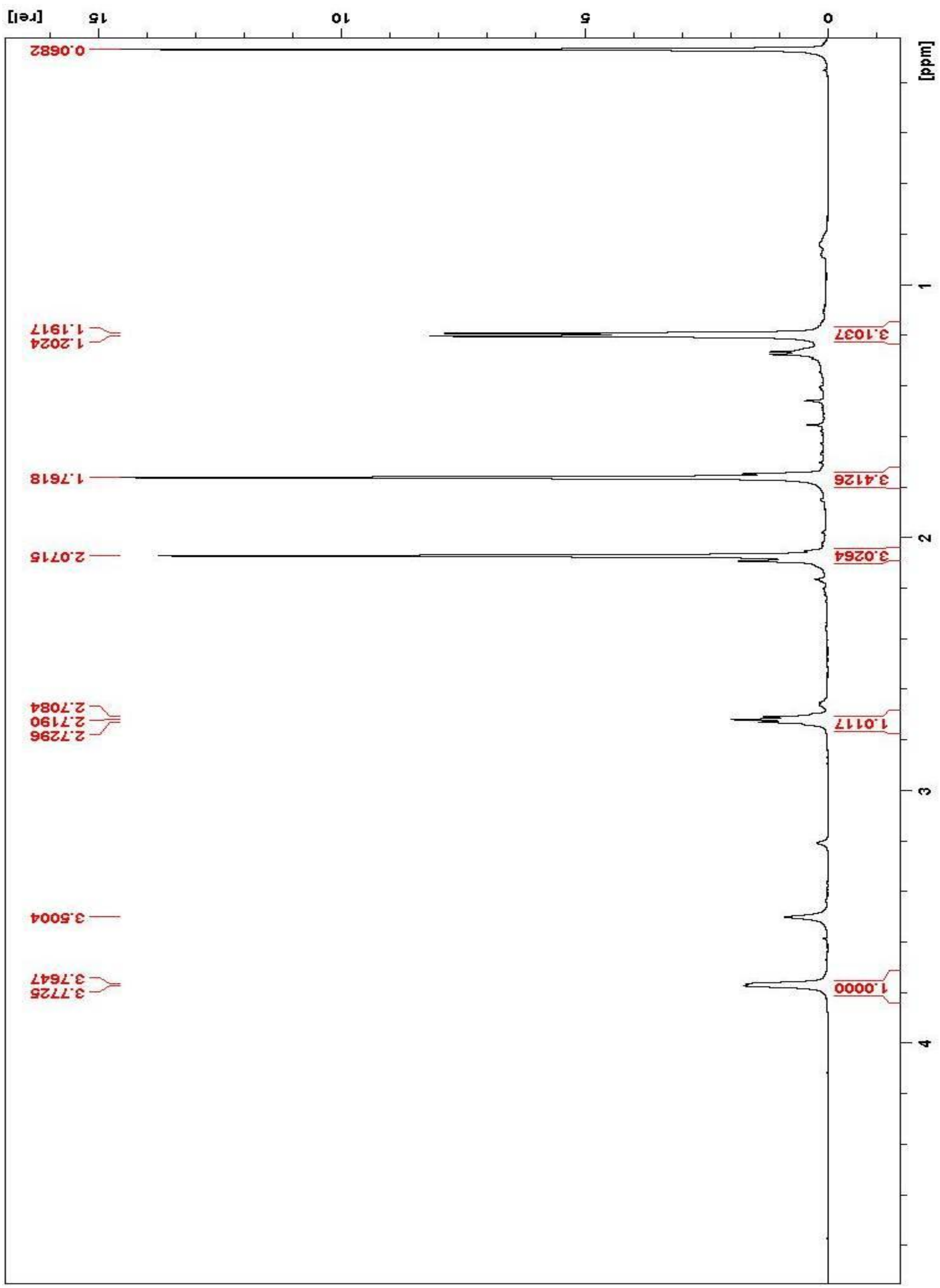


**Figure S21**. $^{1}$H NMR spectrum of methylenomycin D1 (**7**) in $CDCl_3$

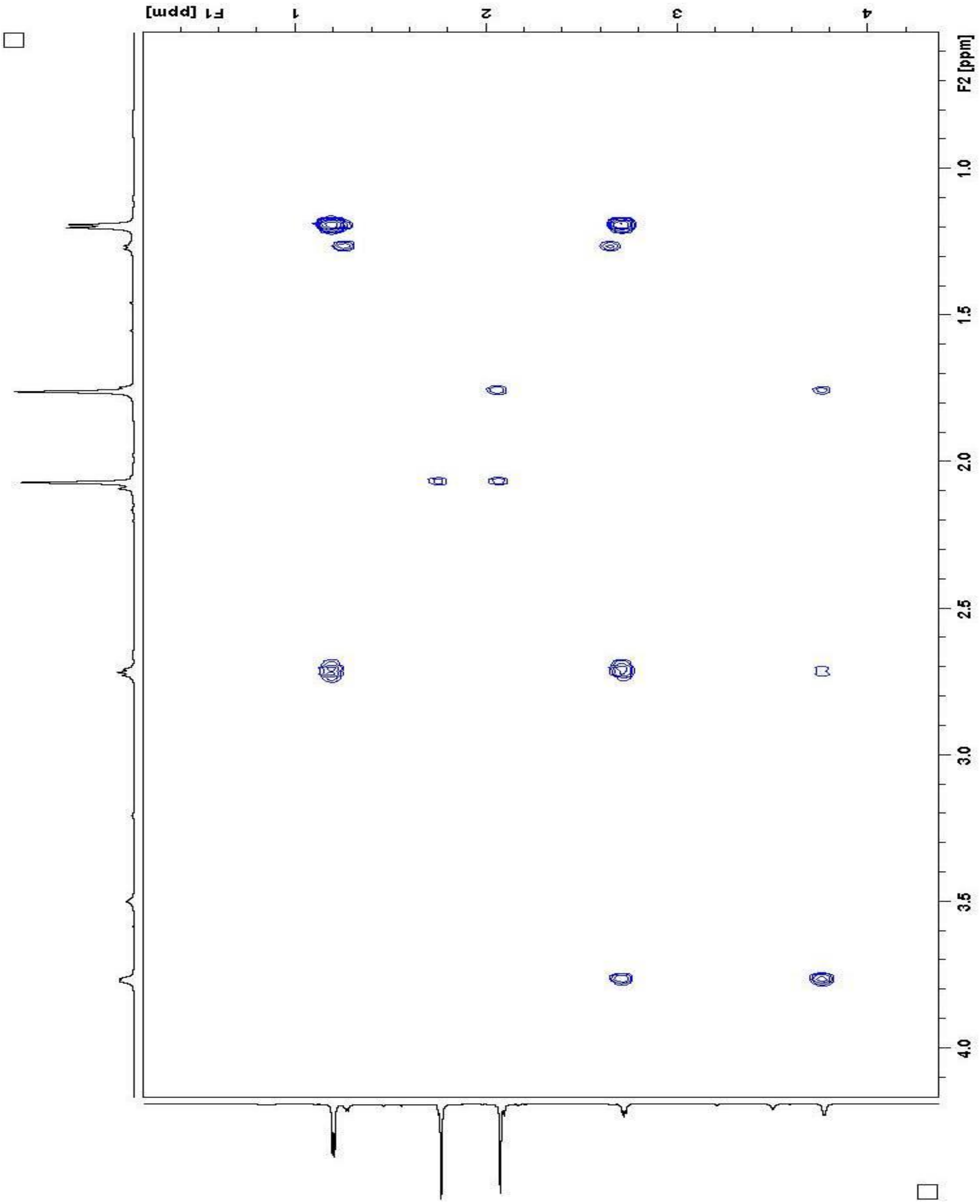


**Figure S22**. COSY spectrum of methylenomycin D1 (**7**) in $CDCl_3$

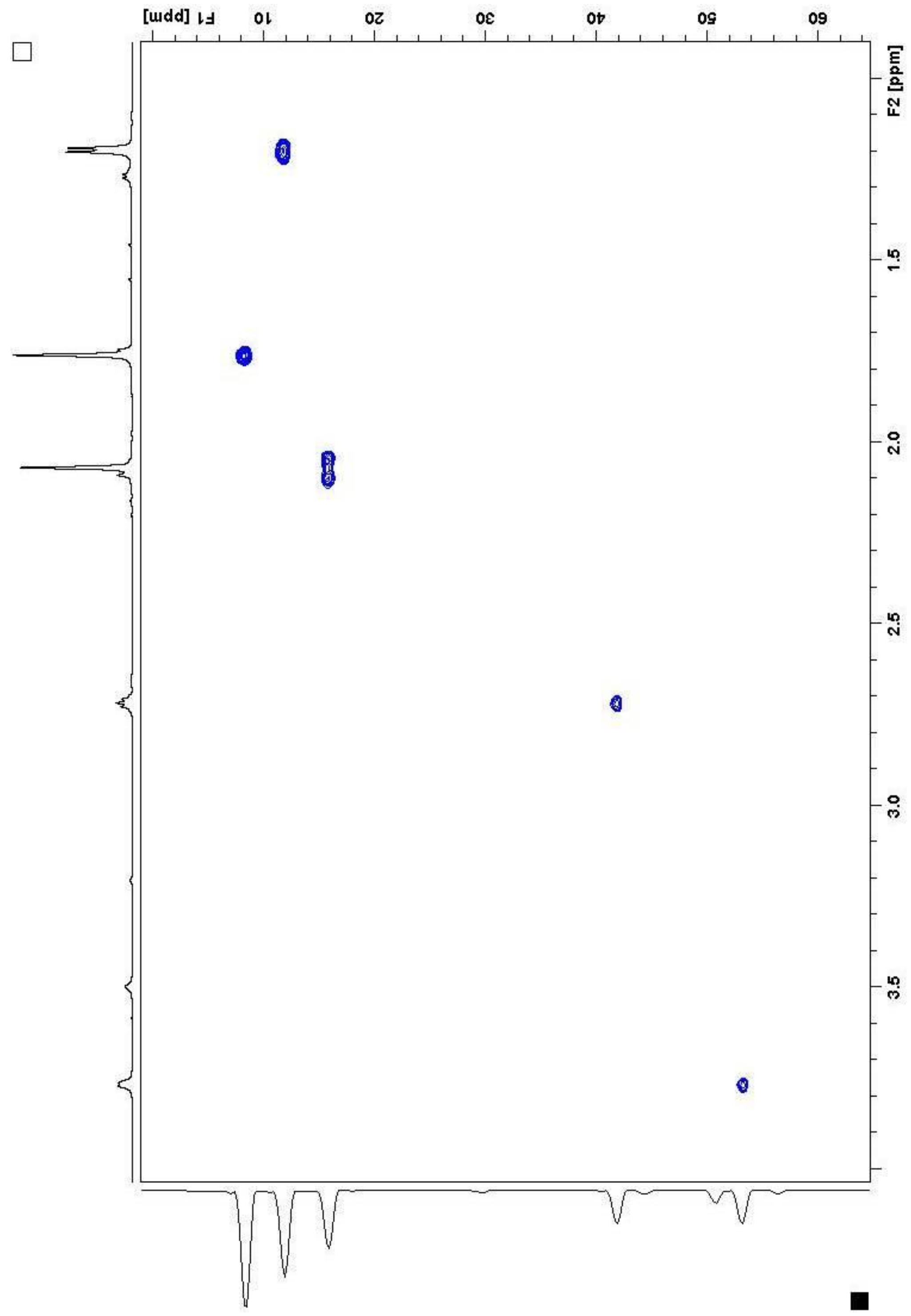


**Figure S23**. HSQC spectrum of methylenomycin D1 (**7**) in $CDCl_3$

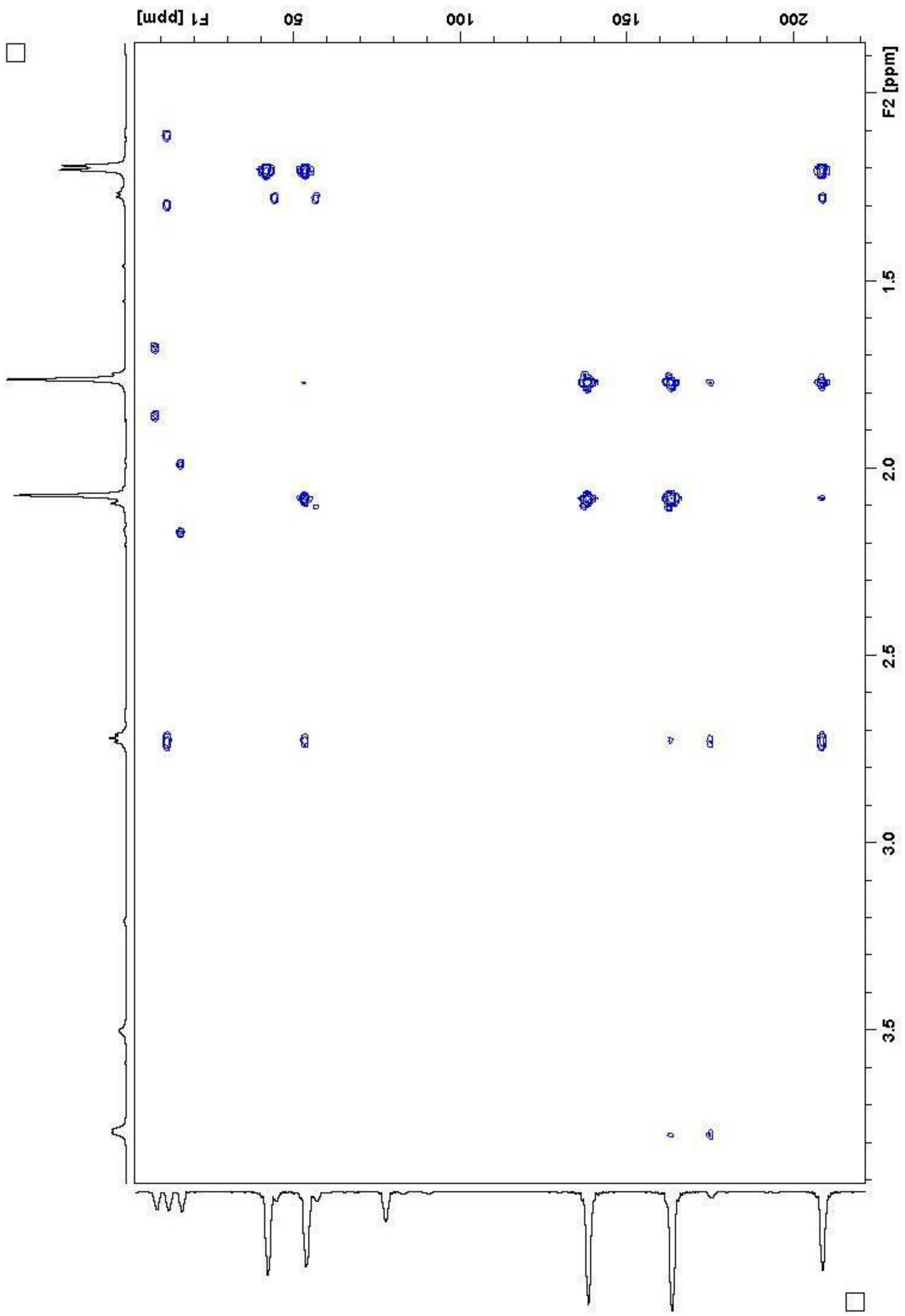


**Figure S24**. HMBC spectrum of methylenomycin D1 (**7**) in $CDCl_3$

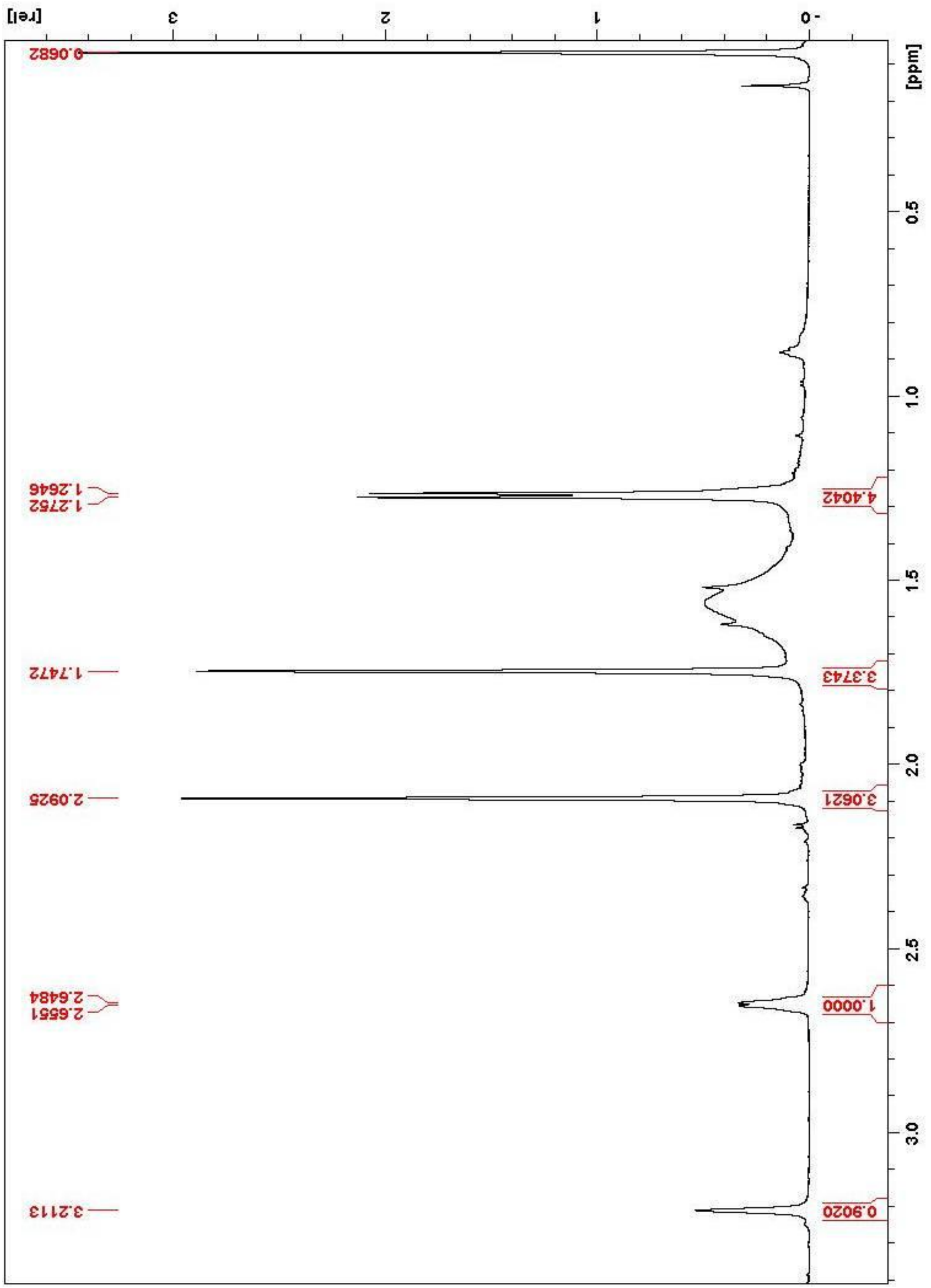


**Figure S25**. [1]H NMR spectrum of methylenomycin D2 (**8**) in $CDCl_3$

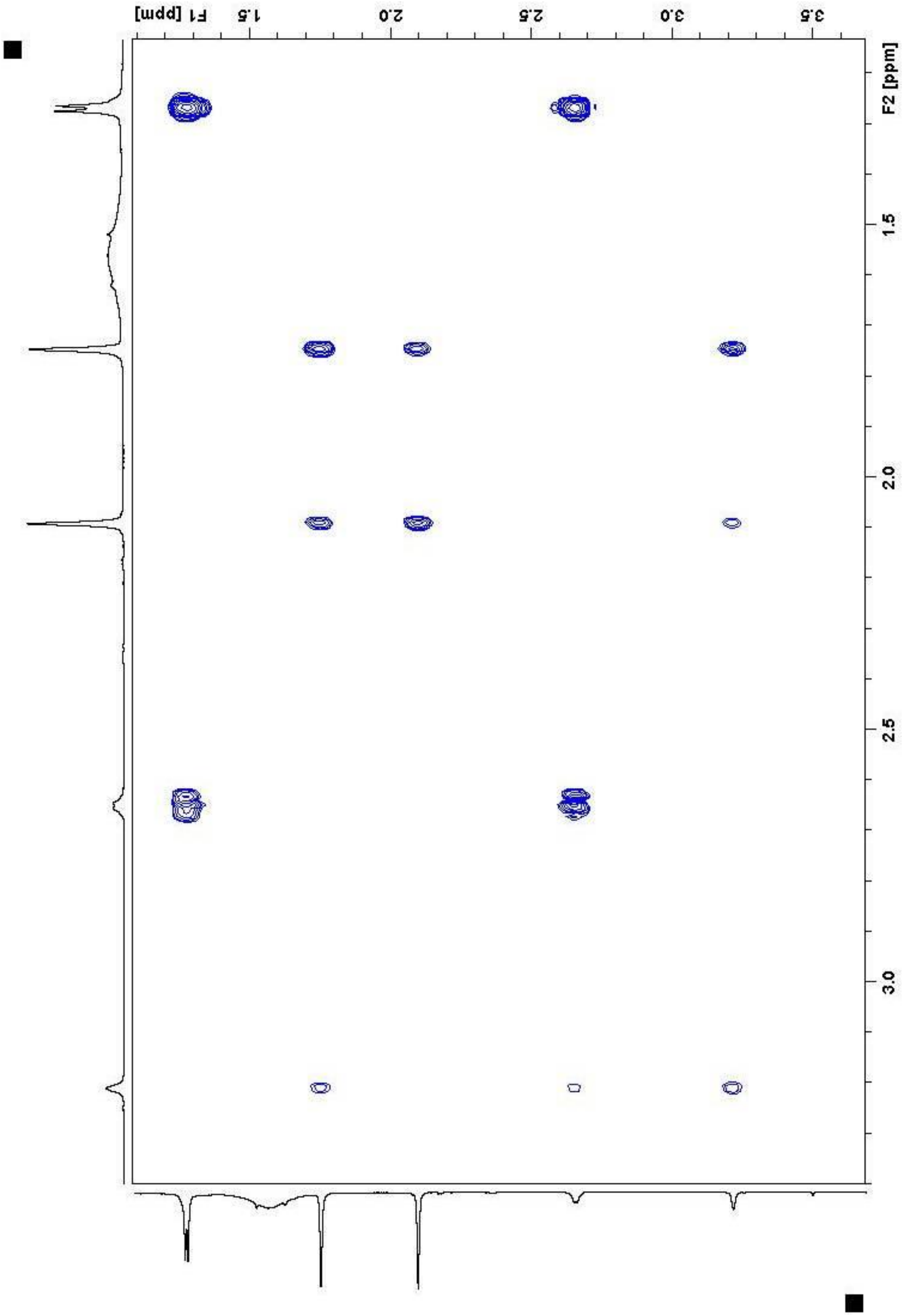


**Figure S26**. COSY spectrum of methylenomycin D2 (**8**) in $CDCl_3$

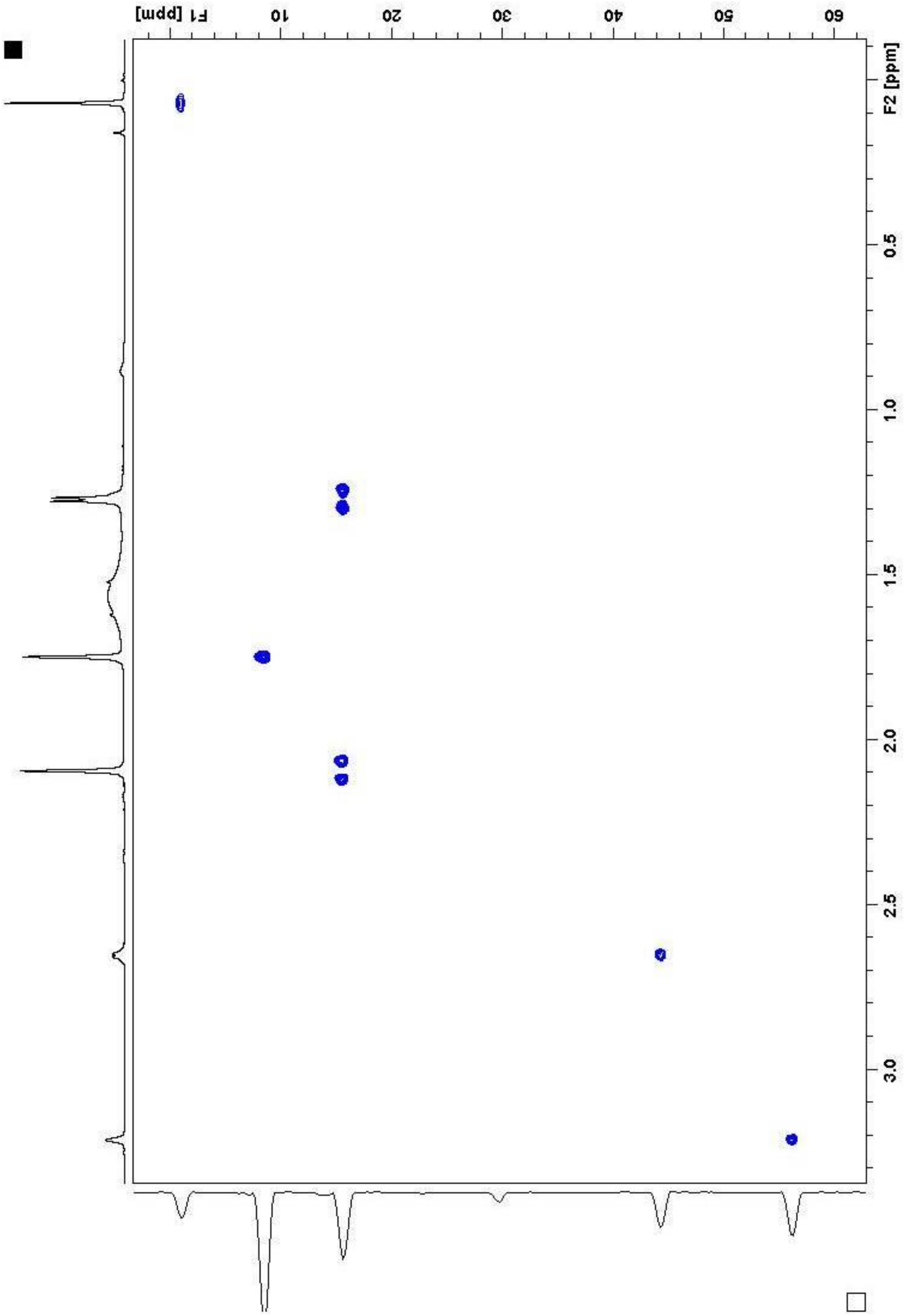


**Figure S27**. HSQC spectrum of methylenomycin D2 (**8**) in $CDCl_3$

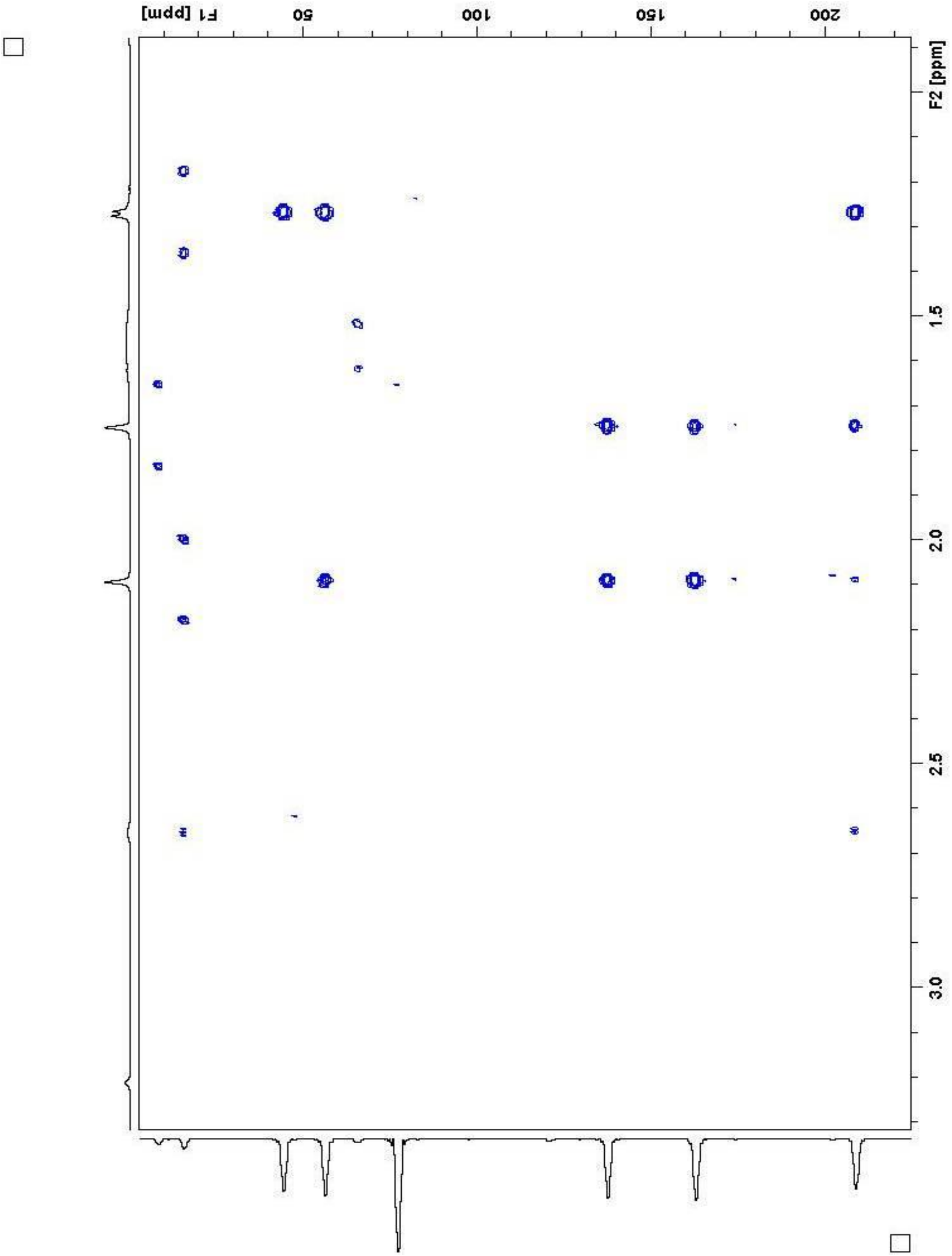

**Figure S28**. HMBC spectrum of methylenomycin D2 (**8**) in $CDCl_3$

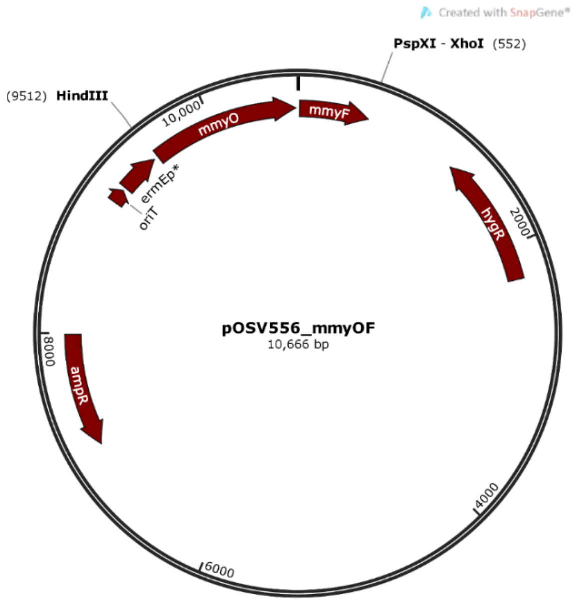


**Figure S29**. Map of pOSV556 carrying *mmyO* and *mmyF*.

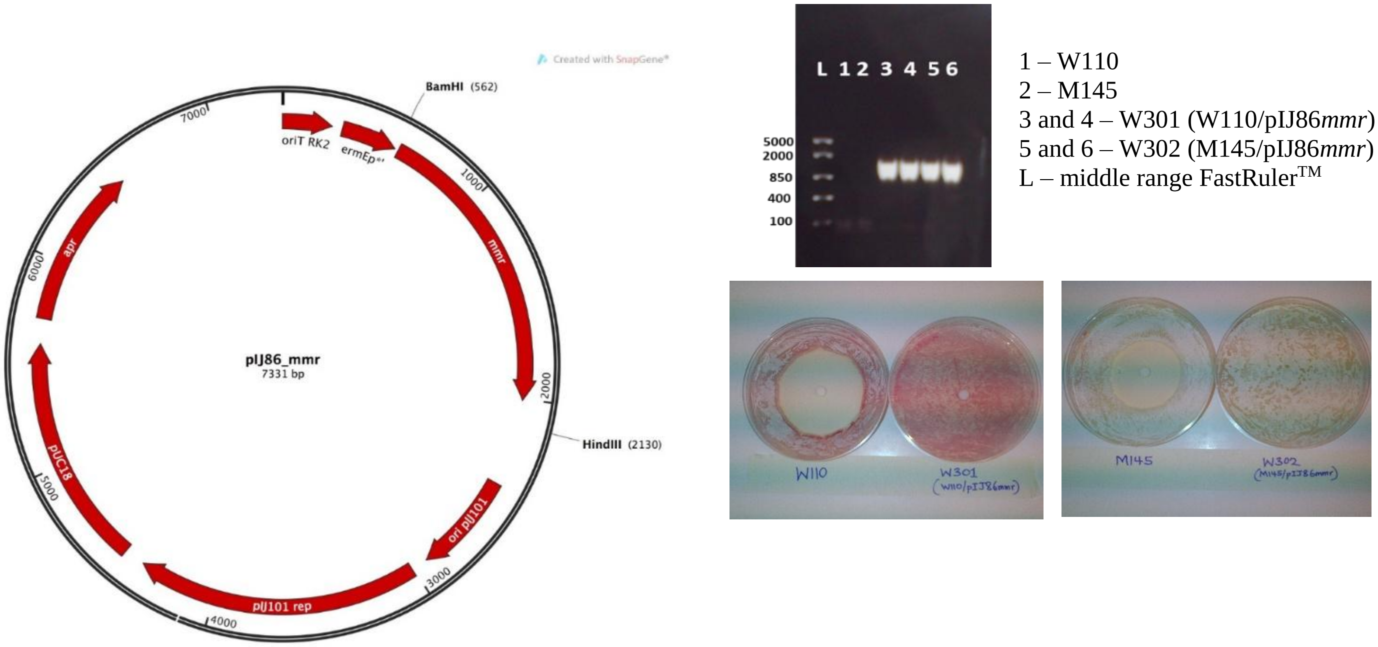


**Figure S30**. Map of pIJ86 carrying the methylenomycin resistance gene, *mmr* (Left). Construction of methylenomycin-resistant strains *of S. coelicolor* (Right): Top is colony PCR confirming presence of the 1.43kb *mmr* DNA in transconjugant strains W301 and W302, and absence from the parent strains W110 and M145, respectively. Filter paper discs impregnated with methylenomycin A caused a zone of inhibition on plates with the parent strains but not in their derivatives harbouring *mmr*.

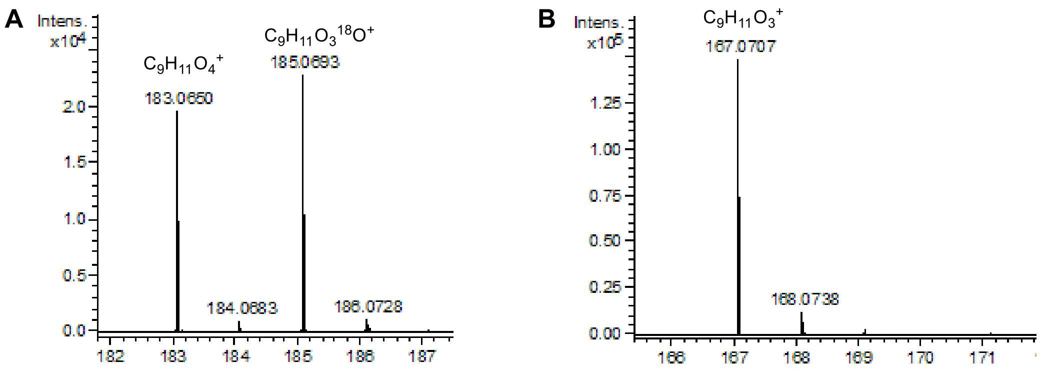


**Figure S31.** High resolution mass spectra of **1** and **2** from UHPLC-ESI-Q-ToF-MS analyses of organic extracts from *S. coelicolor* W89 grown under an $^{18}O_2$ atmosphere. (**A**) The spectrum from the peak corresponding to **1** revealed ions with *m/z* = 183.0650 and 185.0693, corresponding to $[M+H]^+$ for the unlabeled and singly 18O-labelled compound, respectively. (**B**) The spectrum from the peak corresponding to **2** revealed only an ion with *m/z* = 167.0707, corresponding to $[M+H]^+$ for the unlabeled species.

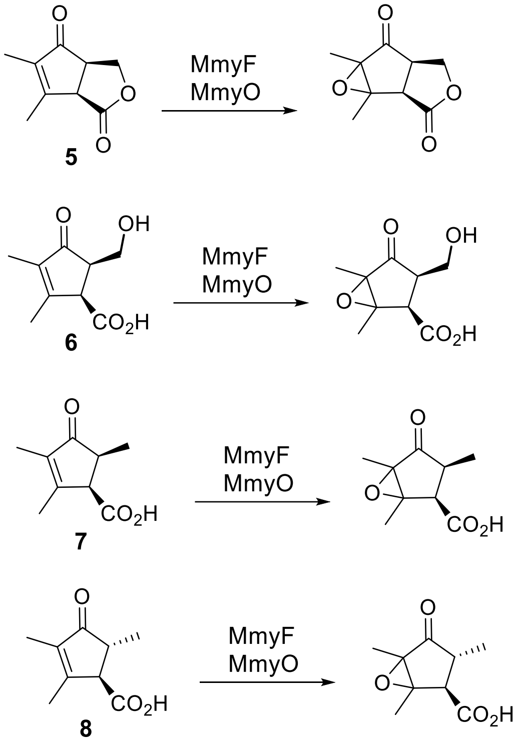


**Figure S32**. Possible products of epoxidation of pre-methylenomycin C lactone (**5**), pre-methylenomycin C (**6**), methylenomycin D1 (**7**) and methylenomycin D2 (**8**) by MmyF and MmyO.

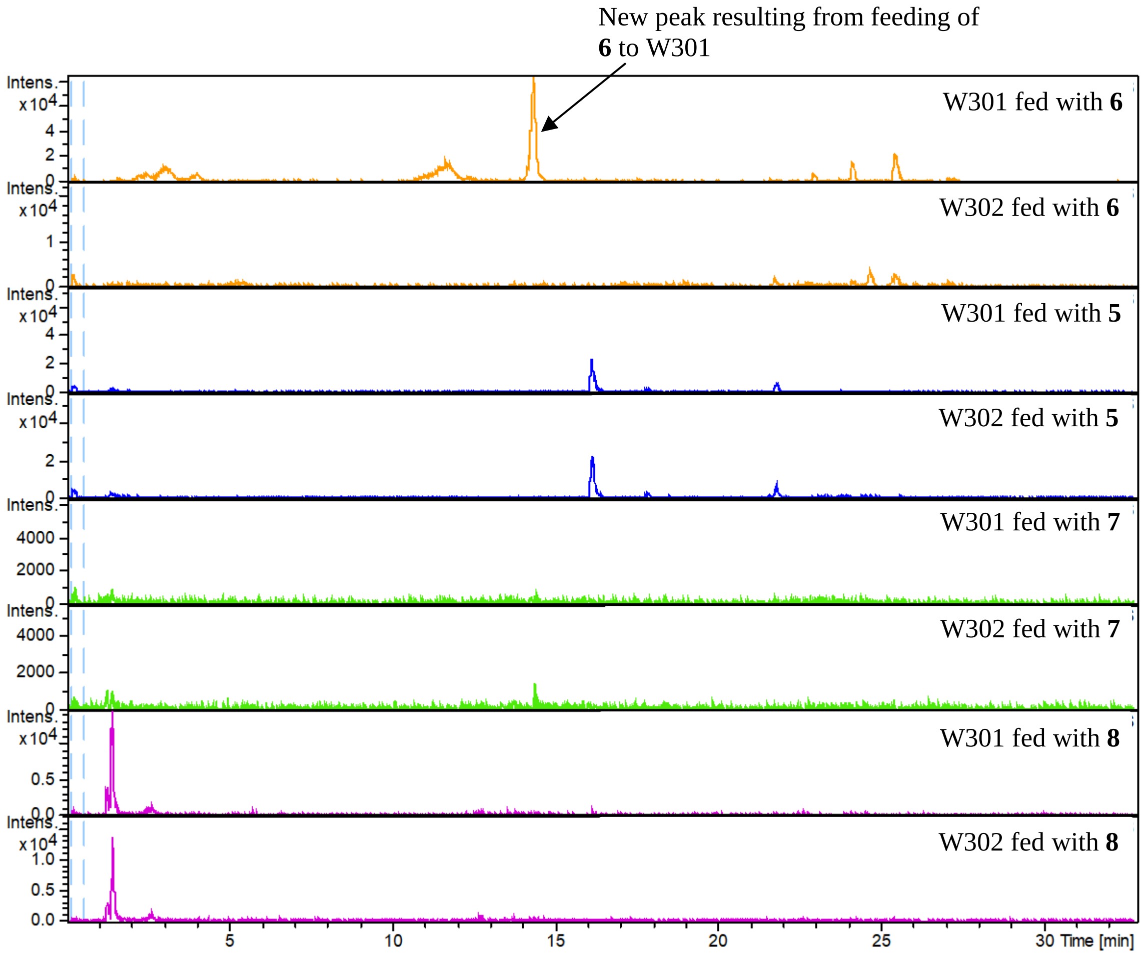


**Figure S33**. Extracted ion chromatograms (EICs) from LC-MS analyses of extracts of *S. coelicolor* W301 and W302 fed with **6**, **5**, **7** and **8**. Yellow: EIC at *m/z* = 201.07 and 223.07 corresponding to $[M + H]^+$ and $[M + Na]^+$ for possible epoxidized product of **6**. Blue: EIC at *m/z* = 183.06 and 205.06 corresponding to $[M + H]^+$ and $[M + Na]^+$ for possible epoxidized product of **5.** Green and Purple: EIC at *m/z* = 185.07 and 207.07 corresponding to $[M + H]^+$ and $[M + Na]^+$ for possible epoxidized products of **7** and **8** respectively**.**

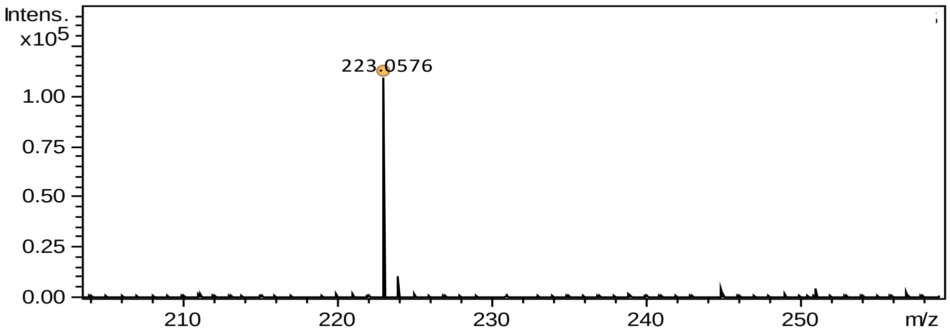


**Figure S34**. Mass spectrum of compound corresponding to the $[M + Na]^+$ ion for the epoxidized product (calculated *m/z* = 223.0577) resulting from the feeding of pre-methylenomycin C (**6**) to *S. coelicolor* W301.

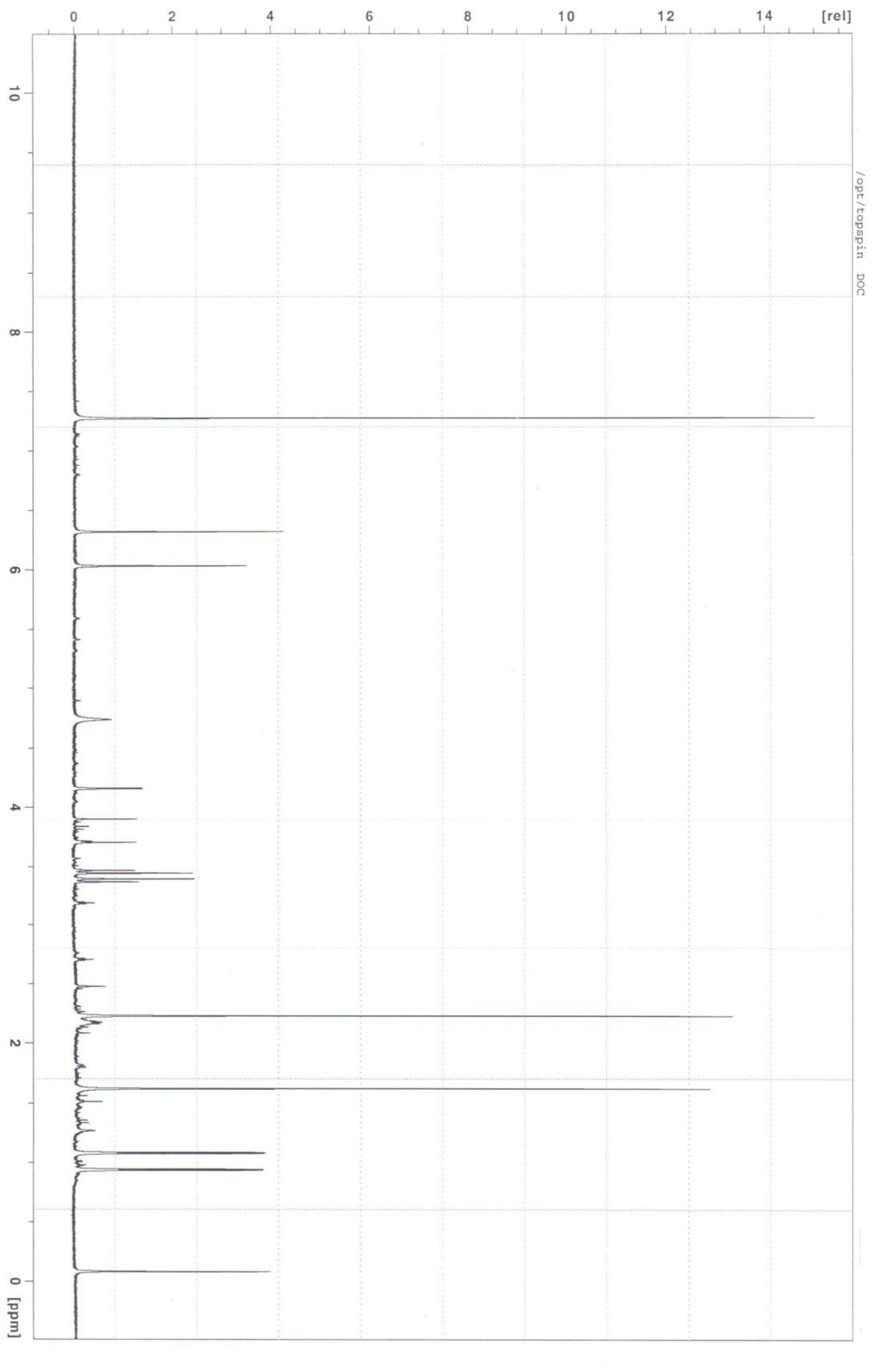


**Figure S35**. $^{1}$H NMR spectrum of **10** in $CDCl_3$

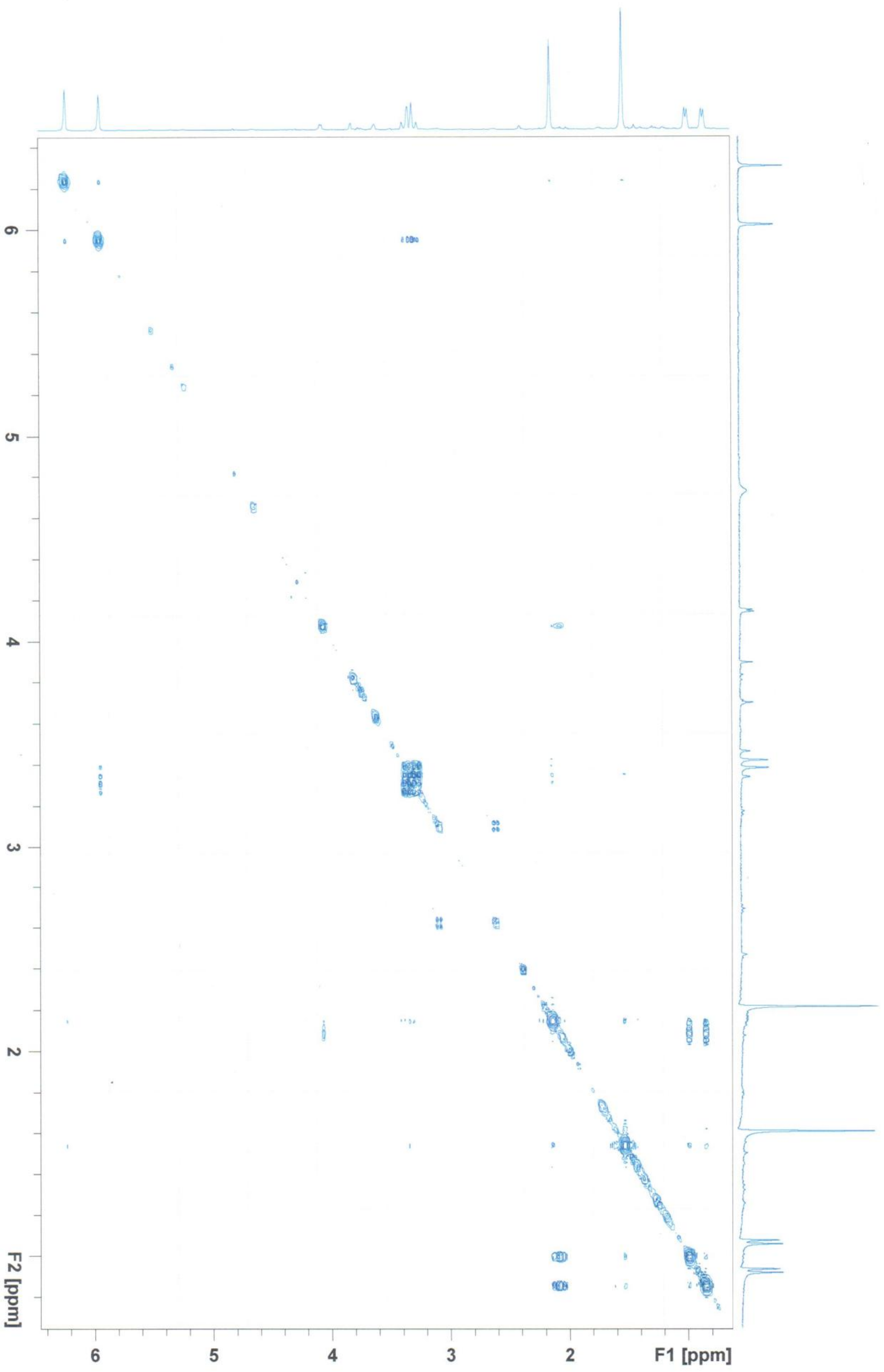


**Figure S36**. COSY spectrum of **10** in $CDCl_3$

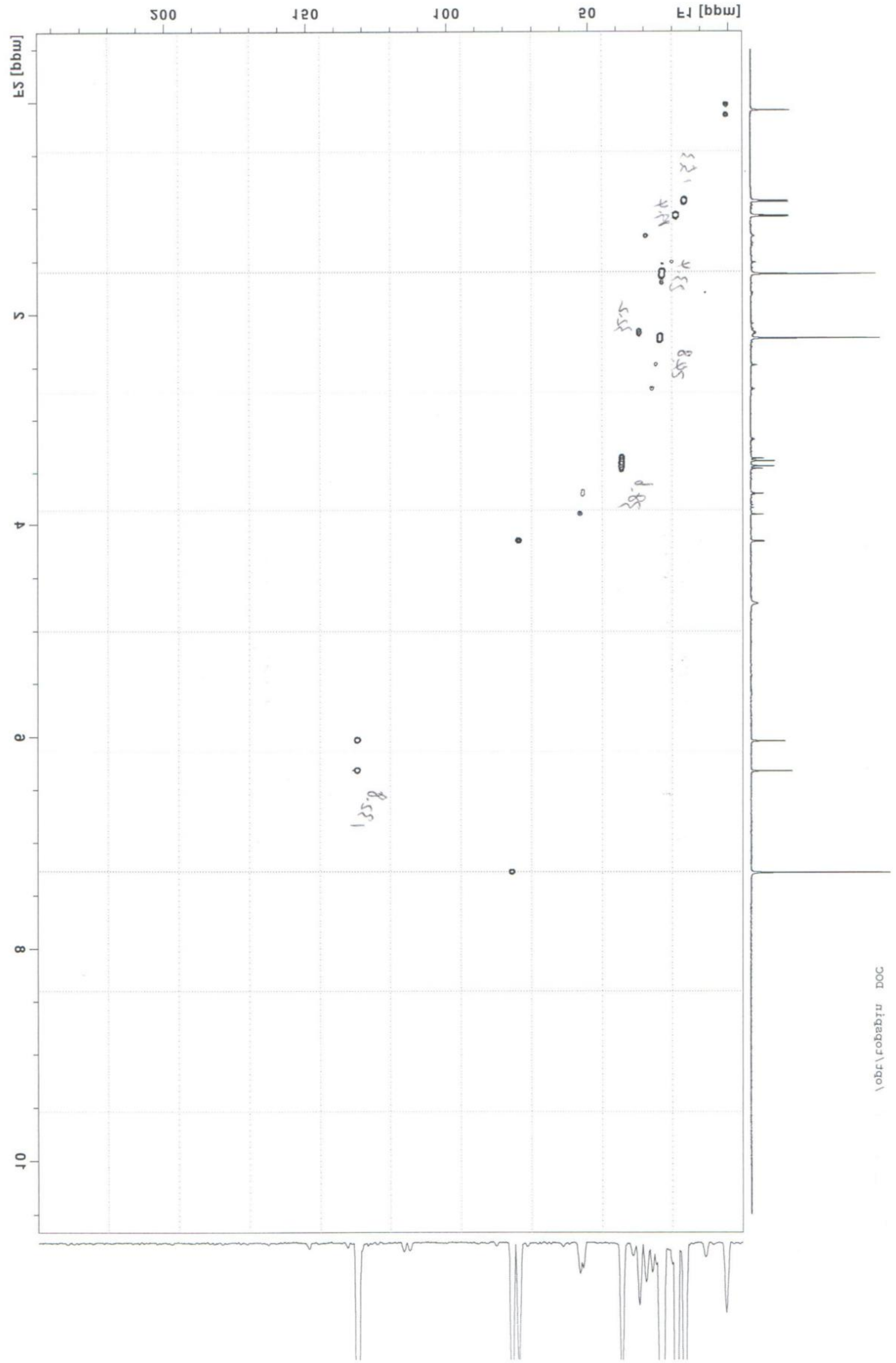

**Figure S37**. HSQC spectrum of **10** in $CDCl_3$

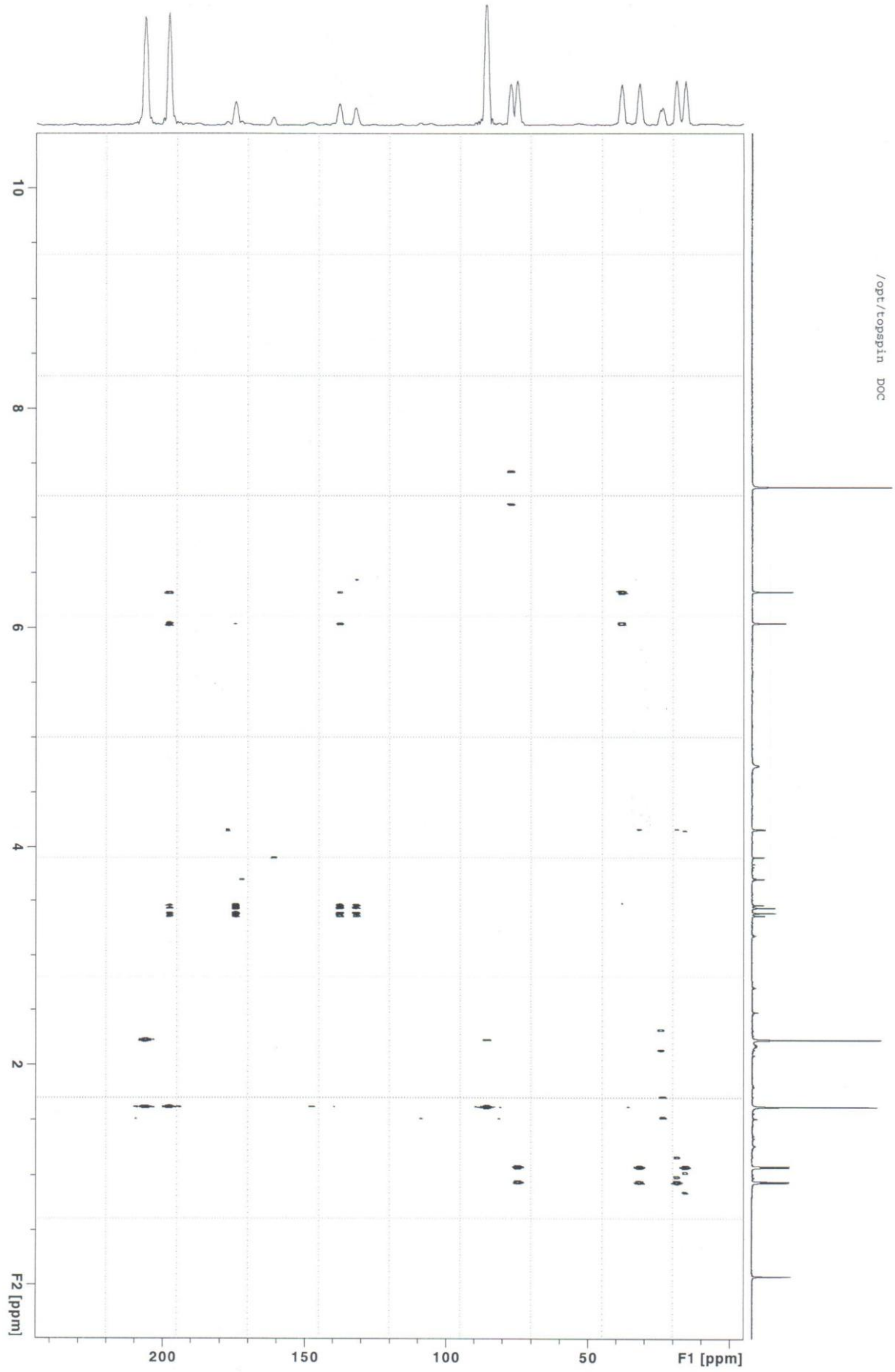


**Figure S38**. HMBC spectrum of **10** in $CDCl_3$
